# Multidimensional cerebellar computations for flexible kinematic control of movements

**DOI:** 10.1101/2022.01.11.475785

**Authors:** Akshay Markanday, Sungho Hong, Junya Inoue, Erik De Schutter, Peter Thier

## Abstract

Both the environment and our body keep changing dynamically. Hence, ensuring movement precision requires adaptation to multiple demands occurring simultaneously. Here we show that the cerebellum performs the necessary multi-dimensional computations for the flexible control of different movement parameters depending on the prevailing context. This conclusion is based on the identification of a manifold-like activity in both mossy fibers (MF, network input) and Purkinje cells (PC, output), recorded from monkeys performing a saccade task. Unlike MFs, the properties of PC manifolds developed selective representations of individual movement parameters. Error feedback-driven climbing fiber input modulated the PC manifolds to predict specific, error type-dependent changes in subsequent actions. Furthermore, a feed-forward network model that simulated MF-to-PC transformations revealed that amplification and restructuring of the lesser variability in the MF activity is a pivotal circuit mechanism. Therefore, flexible control of movement by the cerebellum crucially depends on its capacity for multi-dimensional computations.

## Main text

Short-term motor learning is a specific variant of sensorimotor learning. It provides the ability to rapidly acquire a new control scheme that allows the motor system to cope with the demands of often unexpected or sudden changes in the external environment^1^. Not only external but also internal changes may require fast adjustments. For instance, the motor plant may change due to muscular fatigue slowing movements. Also, boredom and declining motivation, i.e., cognitive fatigue will reduce the speed of movements. If not too extensive, this slowing of movements—the decline of movement “vigor”—may not necessarily degrade endpoint precision as the speed reduction can be compensated by cranking up the overall movement duration, an adjustment of a distinct parameter that requires the cerebellum^2–4^. However, behavioral studies indicate that parametric control by the cerebellum, deployed to swiftly react to external and internal changes, is not confined to a single kinematic parameter like movement duration. Rather, work on goal-directed eye movements as models of cerebellum-based short-term motor learning have established that adaptation to external and internal changes involves adjustments of several kinematic parameters^2,5–11^.

How does the cerebellum coordinate the control of multiple kinematic parameters in order to ensure optimal movements? To answer this question, we should know at which stage of the cerebellar neural network the information on the various movement parameters and necessary adjustments is available and how they are transformed within the network. Previous studies on saccadic eye movements have emphasized the control of particular parameters like movement duration^12^ or velocity^13^ by the simple spike (SS) discharge of a population of cerebellar Purkinje cells (PCs)—the output currency of cerebellar cortex. Although it has been suggested that SS firing rate and spike time can simultaneously encode the velocity and timing of eye movement at the individual PC level^14^, ultimately unifying these divergent views at the population level is challenged by the large cell-to-cell variability of the discharge of cerebellar neurons. This issue is usually addressed by extensive averaging of all or categorized subsets of neurons in data^6,12,13,15,16^. However, averaging can lead to conclusions that are biased towards a particular parameter within a space of multiple encoded movement parameters.

Unraveling the information hidden in cell-to-cell variabilities of neuronal populations is where recent studies of the neural dynamics of cortical motor regions have made remarkable progress^17–20^. One of the key ideas is that the apparent substantial heterogeneity or high dimensionality of the responses of individual neurons can actually be explained by a combination of a smaller number of underlying patterns, i.e., a low-dimensional latent structure. This low-dimensional structure, referred to as the ‘*manifold’*, captures the essential properties residing inside the population discharge^20–23^ without the risk of biased conclusions, inevitably introduced by simple averaging across neurons.

Hence, to address if and how the cerebellum is able to accommodate the multifarious parametric requirements of short-term sensorimotor learning, we identify the manifold structure of the activity of key input and output elements of the cerebellar cortical network, mossy fibers (MFs) and PCs, of nonhuman primates performing a fatigue-inducing repetitive saccade task entailing different kinematic changes. We report the multi-dimensional manifolds in the MF and PC activity that simultaneously encode kinematic parameters, eye movement velocity and duration, by their geometry and dynamics. We then proceed with considering the influence of climbing fibers, represented by the PC complex spike (CS) discharge, conveying information on error feedback, on the PC manifolds. We show that CSs modulate PC manifolds in an error type-dependent manner that predicts complementary changes in subsequent eye movements by selectively controlling the individual movement parameters. Finally, we investigate the nature of the interaction between the input and output neurons and present evidence that the underlying network computation amplifies the relatively small variability in MF responses to transform them into representations of individual movement parameters, exhibited by PCs in an error-type-dependent manner. Our results demonstrate an enhanced computational capacity of PCs that provides the flexible control of more than one kinematic parameter, ensuring the precision of goal-directed movements.

## Results

### Velocity-duration adjustments during a fatigue-inducing repetitive saccade task

We trained two monkeys to execute a long series of visually guided saccades made centrifugally towards a fixed target location (eccentricity: 15 deg) in a horizontally left- or rightward direction in order to receive a water-based reward at the end of the movement (**Fig. 1a**, see methods for details).

**Figure 1.**
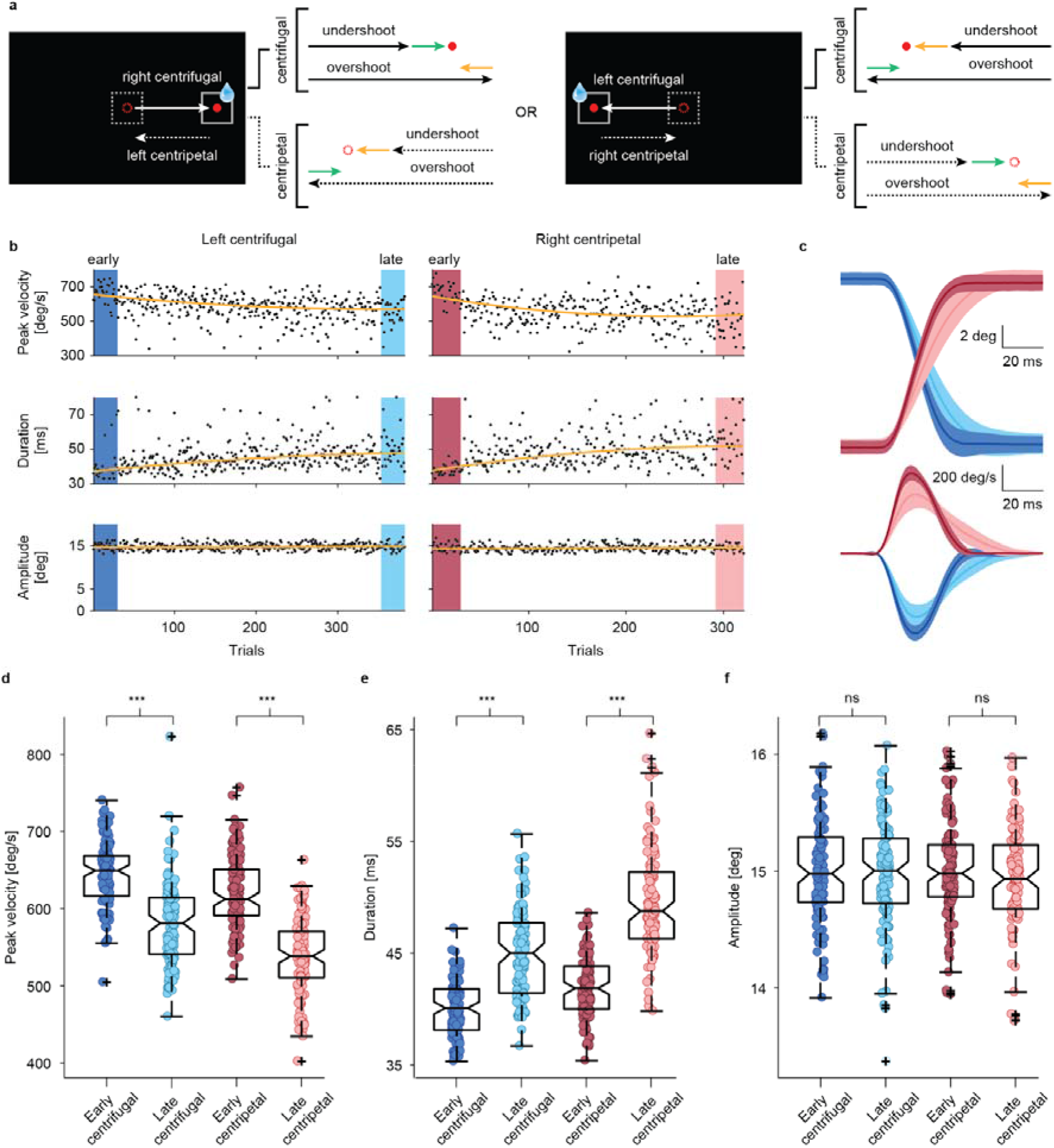
Repetitive saccade task induces a gradual decline in saccade velocity. **a** Behavioral task. Monkeys were trained to make visually guided saccades to targets, either in left or right directions, in a repetitive manner. Solid arrows represent all center-out (centrifugal) saccades which were rewarded if the eyes landed within the 2×2 deg fixation window (solid and dashed squares). Saccades made back to the central fixation dot, centripetal saccades (dashed arrows), were not rewarded. Due to natural variability in eye movements, both centrifugal and centripetal saccades could either overshoot or undershoot the target leading to errors in leftward (orange arrows) or rightward (green arrows) direction. **b** Gradual decay of peak velocity (upper panels) in centrifugal (left) and centripetal (right) saccades (Wilcoxon signed-rank test, centrifugal: p<0.001, Z=4.4; centripetal: p<0.001, Z=4.4) is parallel by an increase in saccade duration (middle panels, Wilcoxon signed-rank test, centrifugal: p<0.001, Z= −3.7; centripetal: p<0.001, Z=−4.8) to stabilize amplitudes (lower panels, Wilcoxon signed-rank test, centrifugal: p=0.89, Z= −0.1; centripetal: p=0.95, Z=0.1) within a single session. Each dot represents data from a single trial. Trends in the data are highlighted by fitting second-order polynomial fits (dark yellow lines) to the data. **c** Comparison of horizontal eye position and velocity profiles of early (i.e., first 30 trials, centrifugal: dark blue; centripetal: dark red) and late (i.e., last 30 trials, centrifugal: light blue; centripetal: light red) trials chosen from the experimental session in c. **d, e, f** Population analysis of 117 behavioral sessions. Box plots showing overall reduction of peak velocity (Wilcoxon signed-rank test, centrifugal: p<0.001, Z=8.6; centripetal: p<0.001, Z=9.3) in late trials (lighter colors) as compared to early (darker colors) ones which is compensated by the upregulation of saccade duration(Wilcoxon signed-rank test, centrifugal & centripetal: p<0.001, Z= −9.3) during the late trials in order to maintain saccade amplitude around 15 deg (Wilcoxon signed-rank test, centrifugal: p=0.57, Z= 0.6; centripetal: p=0.01, Z=2.5). Each data point corresponds to the mean value of the early (first 30, dark-colored circles) and late (last 30, light-colored circles) centrifugal (blue circles) and centripetal saccades (red circles) of an individual session. Significant differences are highlighted by asterisks. Data are mean±SEM.

As exemplified in **Fig. 1 b-d**, saccades exhibited a gradual decline in their peak velocity (PV) over the course of a session, reflecting a general loss of motivation (“cognitive fatigue”), arguably due to the fast and repetitive nature of the task^4^ (**Fig. 1b**, up). This gradual drop in saccade velocities was compensated by a likewise gradual upregulation of saccade duration (**Fig. 1b**, middle) ensuring that endpoint accuracy was maintained (**Fig. 1b**, bottom) within an acceptable range of error (±2 deg around the target). Since inter-trial intervals were short (~100 ms), the monkeys had to execute rapid saccades back towards the fixation point (i.e., centripetal saccades) after every centrifugal saccade to get ready for the subsequent trial. Albeit not directly rewarded, the kinematic structure and the velocity-duration adjustments of centripetal saccades were very similar to those of centrifugal saccades (red and blue traces, **Fig. 1 b,c**). The notion of a viable velocity-duration tradeoff suggested by the exemplary data received full support from a behavioral population analysis which was based on pooled saccades from all sessions in which we had recorded the responses of 117 MFs and complementary dataset of saccades collected while recording from 151 PCs, the latter the basis of Markanday et. al 2021^24^ (**Fig. 1d-f**). Relative to the early trials, we observed an overall decrease of 9.9% in the median PV of late centrifugal and 12.1% decrease in late centripetal saccades (**Fig. 1d**), compensated by a 12.2% and 16.5%, respectively, increases in median saccade duration (**Fig. 1e**), maintaining the required accuracy (**Fig. 1f**).

On top of these gradual changes, reflecting the consequences of the development of cognitive fatigue over many trials, we also observed a within-session, trial-to-trial variability in centrifugal and centripetal saccade endpoints, (‘motor noise’), which resulted from saccades randomly overshooting or undershooting the target (**Fig. 1a**, see schematic diagrams with green and yellow-colored arrows). As a consequence, both saccade types could result in retinal errors in both directions that we could resort to when trying to estimate the preferred error direction of complex spike (CS) firing of individual PCs as projected on the left-right axis.

### Mossy fiber discharge encodes saccade kinematics

We recorded the saccade-related discharge of 117 MFs from the oculomotor vermis(OMV)^25,26^ and broadly categorized them into three main types—burst-tonic (BT), short-lead burst (SLB), and long-lead burst (LLB) units (**Fig. 2b**, see **Materials and Methods** for details) considering the timing of a burst-response component and the presence of subsequent tonic discharge. As demonstrated by an exemplary BT unit (**Fig. 2a**, left panels), a strong “burst” discharge for saccades made in the preferred horizontal direction (here: leftward) direction was followed by an elevated discharge rate (the tonic component), that persisted throughout the post-saccadic period and stopped only when the eyes began to move in the opposite (non-preferred) direction. Compared to the SLB units that started to fire vigorously just a few milliseconds before saccade onset (**Fig. 2a**, middle; 9 ms in the example), the modulation onset of the LLB units occurred much earlier (**Fig. 2a**, right; ~330 ms in the example), reaching its maximum expression in a ramping manner. Independent of MF unit type, the discharge rate reached its peak during the saccade and stopped around the end of saccades, made into a unit’s preferred direction.

**Figure 2.**
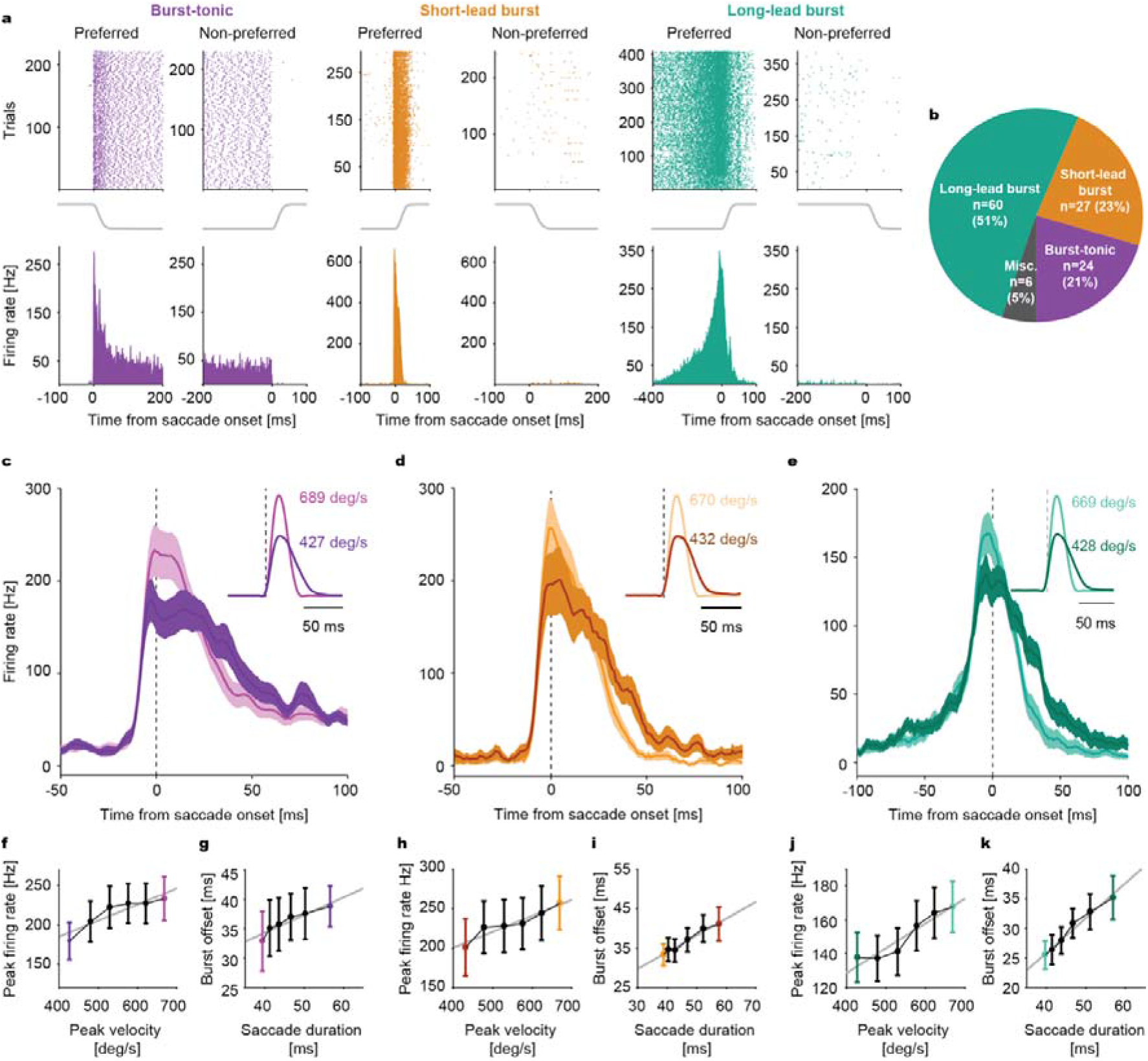
Encoding of saccade kinematics by mossy fibers (MFs). **a** Raster plots (up) and average firing histogram (bottom) of a representative burst-tonic (purple), short-lead burst (yellow) and long-lead burst (turquoise) MF unit. Solid gray lines between upper and lower panels are the mean horizontal eye position traces. Data are aligned to saccade onset. **b** Proportion of MF units in each category. **c, d, e** Population response of burst-tonic (purple), short-lead burst (yellow) and long-lead burst (turquoise) MFs to high and low velocity saccades (see insets for average velocity profiles), represented by lighter and darker shades, respectively. **f, h, j** Average peak firing rate as a function of saccade peak velocity (bin size=50 deg/s) for each MF category. Burst-tonic (f): p=0.016, R^2^=0.83; Short-lead burst (h): p=0.005, R^2^=0.9; Long-lead burst (j): p=0.006, R^2^=0.9. **g, i, k** Average burst offset relative to saccade onset as a function of saccade duration (calculated from velocity bins) for each MF category. Burst-tonic (g): p=0.008, R^2^=0.88; Short-lead burst (i): p=0.0005, R^2^=0.96; Long-lead burst (k): p=0.0005, R^2^=0.97. Solid gray lines represent the linear regression fits. Dark and light-colored bins correspond to the high and low peak velocity bins, respectively, for which population responses in c, d and e are plotted for comparison. Data are mean±SEM.

The discharge of MFs reflected the trial-to-trial changes in saccade kinematics. To demonstrate this relationship, we calculated the population responses for saccades in a unit’s preferred direction, separately for BT, SLB, and LLB MFs (n=24, 27 and 60, respectively, **Fig. 2b**) and sorted them into bins of PV (bin size=50 deg/s), ranging from low to high values (and corresponding changes in saccade duration). Comparing the MF populations responses for the two extreme bins comprising the lowest and highest velocities, respectively, clearly showed that in all three MF groups (**Fig. 2c-e**), the peak firing rate was substantially larger for the high PV bin, associated with clearly shorter burst duration. Note that in all three classes of MF, the peak discharge rate coincided with saccade onset and, moreover, that not only the saccade profiles but also the associated mean discharge profiles were clearly less skewed for the high PV bin. This was due to a shortening of the saccade deceleration phase and a parallel faster decay of the discharge following the discharge peak. In fact, the peak discharge rate grew linearly with PV over the full range of PV bins (**Fig. 2 f, h, j**), whereas the time of burst offset linearly predicted the time of saccade offset (**Fig. 2g,i,k**). Even for the tiniest corrective microsaccades that occurred either during the fixation period or during the post-saccadic period after under or overshooting saccades, we observed the same linear encoding of these kinematic parameters by the activity of the three MF types (**Supplementary fig. 1**).

### Simple spikes of Purkinje cells encode saccade kinematics

We also recorded from 151 OMV PCs and analyzed their simple spike (SS) responses. Whereas MFs exhibited bursting in their preferred saccade direction and little firing in the non-preferred direction, PC SS patterns for the two opposite directions—although often clearly different (see **Materials and Methods**)—did not follow a comparatively simple rule. Therefore, we considered SS responses for centripetal and centrifugal saccades as independent units and classified them into four main categories—burst (n=107), pause (n=99), burst-pause (n=72) and pause-burst types (n=24), using linear discriminant analysis applied on the first two principal components accrued from a principle component analysis (PCA) of the discharge patterns (**Fig. 3a,c**, see **Materials and Methods**). The response of a typical “burst” and a “pause” unit was characterized by a saccade-related increase or decrease in firing rates, whereas “burst-pause” and “pause-burst” units exhibited both types of changes, yet in opposite succession.

**Figure 3.**
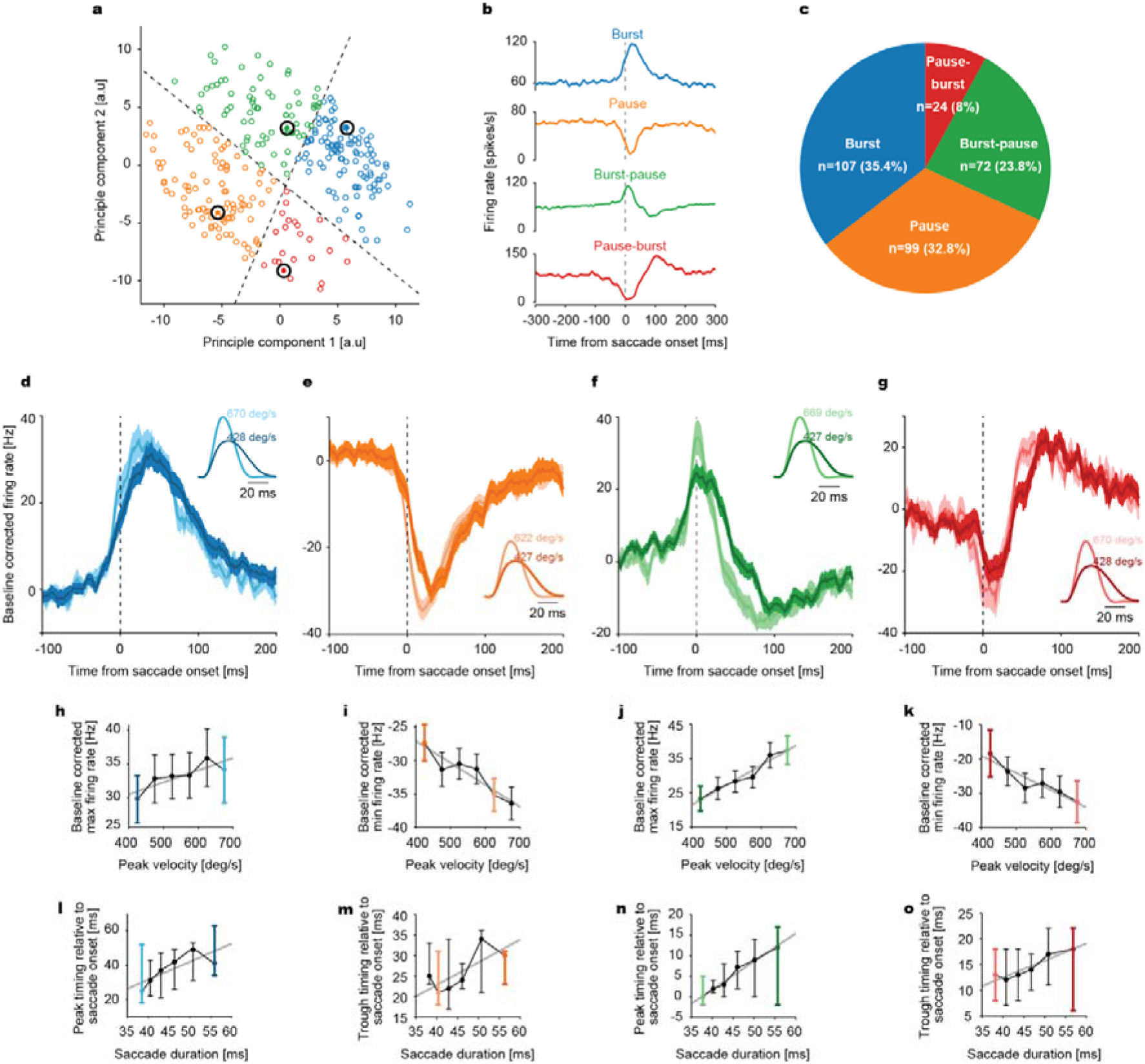
Encoding of saccade kinematics by simple spikes (SSs) of Purkinje cells (PCs). **a** Scatter plot of the first two principal components of SS responses. Classification of PCs into four response categories: burst (blue), pause (orange), burst-pause (green) and pause-burst (red), separated by decision boundaries (dotted black lines). Each data point corresponds to a PC’s SS response in one of the two directions. **b, c** Saccade onset-aligned average SS responses of exemplary units of the categories (large black circles in a) and the proportion of units in each category. **d, e, f, g** SS population response (baseline corrected, mean±SEM) of burst (blue), pause (orange), burst-pause (green) and pause-burst (red) units to high and low velocity saccades (see insets for average velocity profiles), represented by lighter and darker shades, respectively. Data are aligned to saccade onset. **h, i, j, k** Baseline corrected, average maximum (h, j) and minimum (i,k) firing rates as a function of saccade peak velocity (bin size=50 deg/s) for each category. Burst (h): p=0.041, R^2^=0.69; Pause (i): p=0.0065, R^2^=0.87; Burst-pause (j), p=0.00078, R^2^=0.95; Pause-burst (k): p=0.0062, R^2^=0.87. **l, m, n, o** Average peak (for burst and burst-pause units; l, n) and trough (for pause and pause-burst units; m, o) timing relative to saccade onset as a function of saccade duration (calculated from velocity bins) for each PC category. Burst (l): p=0.065, R^2^=0.61; Pause (m): p=0.087, R^2^=0.56; Burst-pause (n): p=0.00015, R^2^=0.98; Pause-burst (o): p=0.0059, R^2^=0.88. Solid gray lines represent the linear regression fits. Dark and light-colored bins correspond to the high and low peak velocity bins, respectively, for which population responses in d, e, f and g are plotted. Data are mean±SEM.

Pooling the responses of all SS units within each category, separately for the aforementioned PV bins, we obtained a clear linear relationship between the firing rate extremes (maximum discharge in units with burst components, minimal discharge in units with pause component) and eye velocity for all four SS categories (**Fig. 3d-g** and **Fig. 3h-k**). To capture saccade duration-related changes in SS firing, we relied on the timing of the first discharge rate extreme. As summarized in **Fig. 3l-o**, it shifted to later times in burst-pause and pause-burst units, while showing the same non-significant tendencies in the other two categories. Hence, one might conclude that the SS discharge of PCs in our data set encoded both movement velocity^13^ and duration^12^.

### Identifying manifolds from pseudo-populations of MFs and PCs to unveil multi-dimensional coding of eye movements

However, there is a necessary caveat. Individual units were recorded in separate sessions. As the trial number varied between sessions and also the behavioral state of a monkey would hardly have been constant over sessions, we cannot exclude that particular velocity bins in our analysis might have been biased by particular sessions, associated with distinct states. Therefore, testing the influence of PV at the population level might be confounded by a potential influence of behavioral state variables, such as movement vigor left uncontrolled. In order to circumvent this potential confound of kinematic dependencies of MF and SSs of PC units in our analysis, we resorted to a computational model that predicted the firing rate of individual MFs and PC-SSs based on a linear combination of a kinematics-independent component, namely the mean firing rate of a unit, and a PV- and/or duration-based modulation as added kinematics-dependent components (see **Materials and Methods**, Equation 1). Finally, by combining the linear models of individual units, we obtained a “pseudo-population” of MFs and PCs (for illustration, using only PV as the kinematic-dependent parameter, see **Supplementary fig. 3** and **Materials and Methods** for more details) in which virtually every unit’s contribution to the population response was equal for any given PV bin as if all units had been recorded simultaneously during an experimental session^22^.

The population responses computed from the pseudo-population model of MFs predicted the actual peak firing rate and duration of the population burst discharge with high accuracy (**Supplementary fig. 3e**). On the other hand, the pseudo-population model for PC-SSs also predicted the same quantities significantly, but less well, in particular the burst duration (**Supplementary fig. 3g**). Note that the quality of the prediction did not improve substantially by considering both parameters (i.e., PV and duration) or only PV (see **Supplementary fig 3e,g** and **supplementary methods**). This is expected, since, for maintenance of endpoint precision, a change of one kinematic parameter must be compensated by a coupled change of the other. Therefore, in most cases, we used the PV-only model to probe the effects of the compensatory duration change correlated to PV change as in **Fig. 2,3**. However, the PV and duration-based models were useful for investigating the effects of the residual, uncorrelated changes in PV and duration (see below). The relatively poor prediction provided by the pseudo-population of PC-SSs might be affected by a much larger variability of the kinematics predictions of the individual models, reflected in higher standard errors of population averages of kinematics-independent and kinematics-dependent components (**Supplementary fig. 3c**, bottom panels). A possible source of the high unit-to-unit variability could be the mixing of SS responses of individual PCs, each preferring a specific direction of retinal error. In fact, it has been shown that the conventional saccade-related SS population averages exhibit higher firing rates if the saccades considered are made in a direction that is opposite to the preferred direction of CSs, the latter the direction associated with the highest probability of observing CSs (CS-ON direction)^13^. Hence, could the performance of the PC-SS pseudo-population kinematics prediction be improved by grouping individual PC-SS responses into two pools that share the preference for error direction, i.e., left and right error, respectively? Indeed, reorganizing our PC data based on CS error-tuning, approximated by deciding whether left- or rightward errors evoked larger CS firing rates, led to a clearer saccade-related burst around the time of the saccade in the CS-OFF direction, whose peak clearly modulated with PV (**Supplementary fig. 4a,b**), unlike for saccades made in the CS-ON direction (**Supplementary fig. 4c,d**). However, despite controlling for preferred error directions leading to qualitative differences in the CS-OFF direction, the performance of the SS model in predicting the actual firing rates and burst duration did not improve, as compared to the performance of the SS pseudo-population response (**Supplementary fig. 3e**) obtained by ignoring the CS-ON and CS-OFF directions, possibly due to prevailing large heterogeneity in SS responses. This is further supported by the results of the PCA of PV-dependent components, where a large number of dimensions were required in the case of PCs (d=10), as compared to MFs (d=4), to explain ~78% of the total cell-to-cell variability (**Supplementary fig. 4e,f**).

In an attempt to mitigate the impact of this apparent large cell-to-cell variability, we referred to the dimensionally reduced representations of the pseudo-population responses of MFs and PC-SSs. To this end, we first ran a PCA on the movement parameter-independent components of the individual MFs and PC-SS firing rate predictions provided by the model to identify the number of dimensions explaining a majority of the total cell-to-cell variability. Then, we computed how each of these dimensions encodes movement parameters (PV or/and duration) using the matrix perturbation theory (see **Methods**, Equation 2). For MFs, in the first step, we found two dimensions that explained 87.6% of the total cell-to-cell variability (**Supplementary fig. 5b**), where the first dimension represented a burst modulation (**Supplementary fig. 5c, top**), similar to the population average firing whose burst size and duration were modulated by PV. The second dimension (**Supplementary fig. 5c, bottom**) represented changes in firing rate that varied more slowly before and after the burst response (observed in Dimension 1) in a biphasic manner, indicating an anti-correlation between the pre-and post-burst firing. However, in PCs, capturing 92.3% of the total cell-to-cell variability required four dimensions, where the first two dimensions represented simple monophasic (i.e., bursting or pausing) and biphasic (burst-pause or pause-burst) firing patterns, respectively, whereas the remaining two dimensions exhibited more complex features (**Supplementary fig. 5f,g**).

Plotting these reduced dimensions as a function of each other, we identified the 2D manifolds of the pseudo-population of MFs and PCs for different values of PV. While both MF and PC manifolds appeared as limit cycle-like rotating trajectories, they exhibited crucial differences from each other (**Supplementary fig. 5d,h** and **Fig. 4**). For example, unlike the MF manifolds that were characterized by an overall PV-related increase in their size almost symmetrically around the saccade onsets, the PC-SS manifolds based on the first two dimensions showed no significant changes before saccade onsets, as depicted by the strong overlapping of the manifolds (**Fig. 4a-e**). However, the PC manifolds for the third and fourth dimensions showed clear differences already before saccade onsets. Therefore, PC manifolds based on different dimensions can selectively encode specific phases of a movement, preparation and execution, in the same manner as the “null-space” in cortical manifolds for the preparation of reaching arm movements^18,19^, while the MF manifolds lacked this information suitable to control specific movement phases.

**Figure 4.**
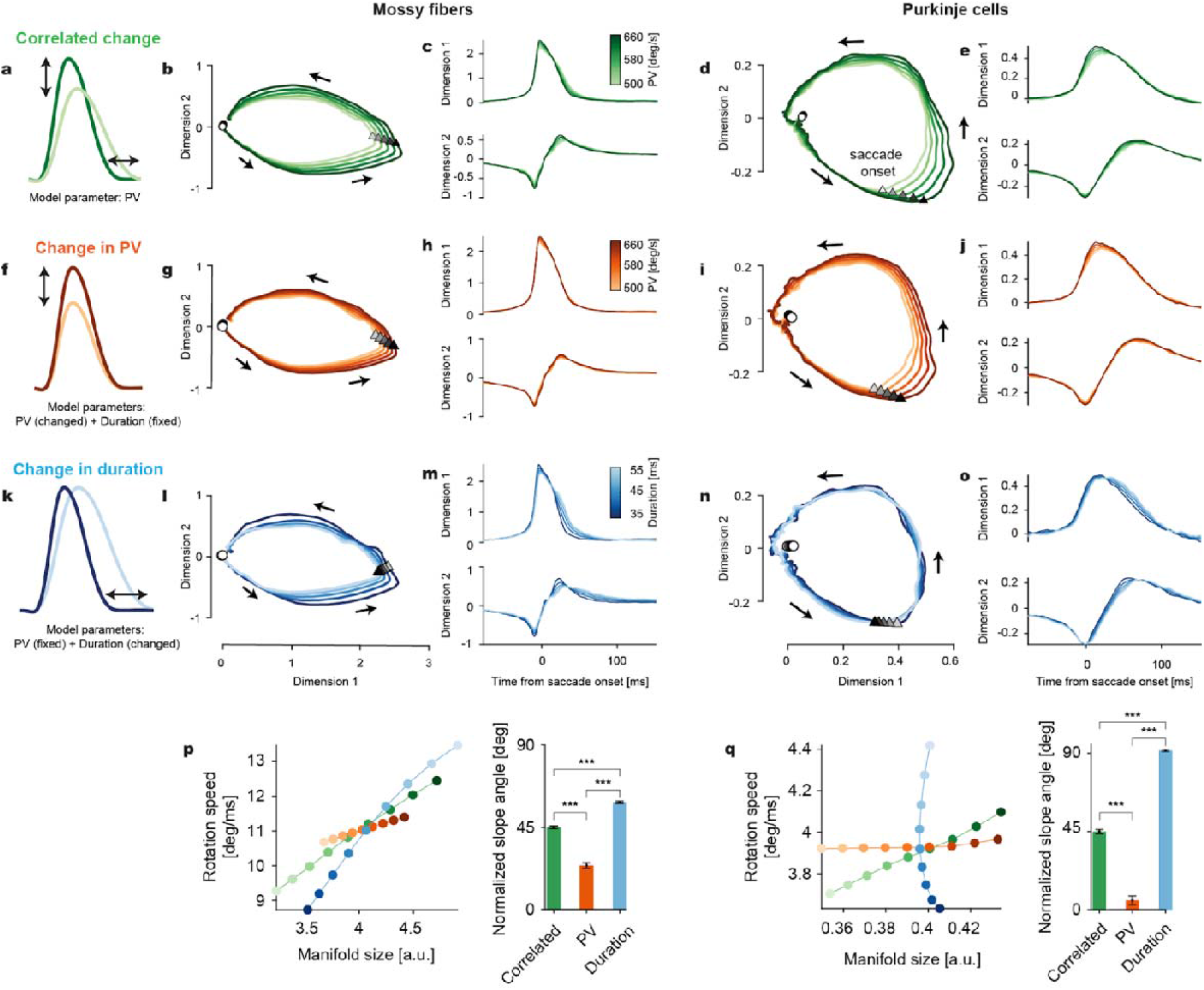
Manifolds identified in MF and PC-SS activity perform multi-dimensional encoding of eye movements. **a** Correlated changes in PV and duration when PV is used as the only control parameter. **b** 2D plot of the first two dimensions in the MF manifold. Triangles and circles mark the saccade onsets and 250 ms before saccade onsets, respectively. Arrows show the direction of rotation. **c** The first two dimensions in b plotted in time. **d,e** Same as b,c for PCs. **f** Isolated changes in saccade PV with the duration kept constant. **g,h** Isolated PV-dependent changes in the MF manifold computed from the rate models parametrized by PV but with fixed duration. **i,j** Same as g,h for PCs. **k** Isolated changes in saccade duration with constant PV. **l-o** Same as g,h and i,j for duration change. **p** Left: MF manifold size versus rotation speed along the MF manifold varying with the correlated (green; a) and independent (orange and blue; f,k) change of PV and duration. Colors are as the color bars in c,h,m. Right: Slope angle of the lines in Left. In computing the angles, the x- and y-coordinates (manifold size and rotation speed) are normalized by the standard deviation of the correlated change case. T-val (Correlated, PV) =17.97; p=1.27×10^−35^, T-val (PV, Duration) =−30.37; p=2.44×10^−57^, T-val (Correlated vs Duration) =−19.18; p=4.46×10^−38^. **q** Same as l for PCs. T-val (Correlated, PV) =19.75; p=5.26×10^−44^, T-val (PV, Duration) =−47.18; p=1.36×10^−92^, T-val (Correlated vs Duration) =−48.13; p=8.24×10^−94^. Data are mean±SEM.

Furthermore, PC manifolds also carried a more disentangled representation of the two saccade parameters— PV and duration, as compared to MFs. To arrive at this conclusion, we estimated MF and PC-SS firing rate models based on both PV and duration by leveraging the residual variabilities in these parameters, apart from their correlated ones. Then, we independently manipulated these two kinematic parameters (varying one while keeping the other fixed) and observed concomitant changes in MF manifolds—a change in PV (**Fig. 4f**) modulated the manifold size (i.e., geometry) and also the time-dependence (i.e., rotation dynamics), the latter still reflecting a small remanence of a correlated duration change (**Fig. 4g,h**). Manipulating the saccade duration (**Fig. 4k**) also modified the MF manifolds (**Fig. 4l,m**) in a manner quite similar to the one resulting from correlated changes in PV and duration (**Fig. 4a**), with PV being the only kinematic parameter in the firing rate model. In contrast, PV (**Fig. 4i,j**) and saccade duration (**Fig. 4n,o**) varied the PC manifold size and rotation dynamics quite differently. These effects are captured by the slope angles of curves obtained by plotting the average rotation speed as a function of manifold size. Therefore, while the slope angles did not differ much in the case of MFs (**Fig. 4p**), the differences were much stronger in the case of PCs, indicating significantly more decorrelated encoding of the two kinematic parameters than MFs (**Fig. 5q**).

**Figure 5.**
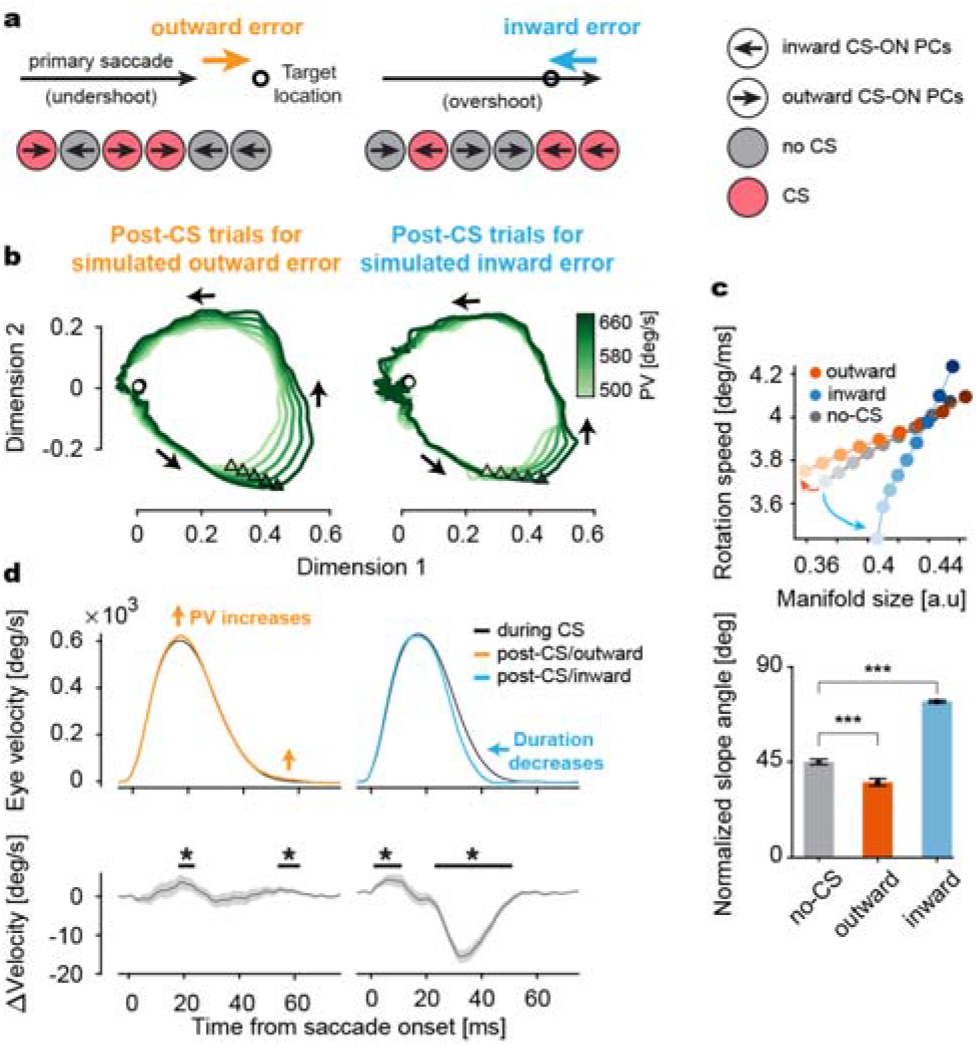
Complex spike (CS)-driven plasticity of PC manifolds is error-state dependent and predicts eye movement change. **a** Two different types of eye movement errors and CS firing in PCs encoding the errors. Left: An undershooting eye movement causes an outward error (orange) and CS firing in a population of PCs with the same CS-ON direction (red) but not in the other, CS-OFF PCs (grey). Right: Same as Left for an overshooting saccade causing an inward error (cyan). **b** Left: PC manifolds reflecting the combined influence of outward error-encoding CS firing pattern in PCs (red and grey circles in a, Left) on subsequent trials. Note that, in simulating error trials by including all CS trials from CS-ON PCs, we assume that PCs are reported an error by CS firing, irrespective of the actual presence of an error. Right: Same as Left for the inward error. **c** Top: Manifold size versus rotation speed after the outward (red) and inward (blue) error-encoding CS-trials, and after no-CS trials (grey). Brightness represents PV from 500 deg/s (brightest) to 660 deg/s (darkest). Bottom: Comparison of normalized slope angles for each condition. T-val (No-CS, Outward) =6.38; p=9.96×10^−10^, T-val (Outward, Inward)=−29.13; p=3.72×10^−64^, T-val (No-CS, Inward)=−28.06; p=4.24×10^−62^. **e** Top: Average saccade velocity profiles in the CS (black) and post-CS trials (colored) for the simulated inward (Left) and outward (Right) errors. For visual clarity, colored lines represent the effect of five CSs in CS-ON cells. Bottom: Average eye velocity change from the CS to post-CS trials. Data are mean±SEM. *: p<0.05.

Centrifugal saccades could be either leftwards or rightwards, but, notably, we found that our results did not depend on saccade direction. To test the potential influence of saccade direction on MF and PC manifolds, we performed the same analysis on MF and PC data separated by leftward and rightward saccades. For MFs, the left and right groups showed qualitatively identical results (**Supplementary fig. 6a,b**). The canonical correlation analysis (CCA)^23,27^ yielded high canonical correlations between the MF manifolds for leftward and rightward saccades, proving that they were nearly identical (**Supplementary fig. 6c**). In the PC case, the size of the manifold was much larger for saccades in the rightward direction as compared to leftward saccades (**Supplementary fig. 6d,e**). Since around 80% of the recorded PCs had their CS-OFF in the rightward direction, the direction-dependent differences in the size of these manifolds are not surprising and only confirm the gain-field encoding of SSs^13^ (**Supplementary fig. 6g,h,i**). Nevertheless, the shape of these manifolds was highly similar (**Supplementary fig. 6f**). Therefore, MFs and PCs had qualitatively identical manifold structures regardless of the eye movement direction.

### PC manifolds reveal the structure of plasticity triggered by sensorimotor errors

In the prevailing theory for cerebellum-dependent sensorimotor learning, the climbing fiber-driven CSs convey motor error-related information to prompt parametric adjustments for correcting future motor behavior, thereby acting as “teacher signals"^28–30^. Therefore, motor learning has been attributed to these CSs, serving as a proxy of sensory feedback on motor errors that, when coincident with the parallel fiber inputs, modify the PC output by inducing a long-term depression (LTD) at the parallel fiber-PC synapses^31^.

To understand how the occurrence of CS impacts the multi-dimensional encoding of eye movements, we investigated how CSs fired during the post-saccadic period of 50-175 ms in the n^th^ trial (‘CS-trial’), reflecting retinal errors arising from natural end-point variability in saccades^24^, modulated the PC-SS manifolds of the subsequent, n+1^th^ trials (‘Post-CS trial’). In our paradigm, errors occurred mainly when the primary saccade undershot (outward error) or overshot (inward error) the target location (**Fig. 5a**). Depending on the direction of the primary saccade, these inward and outward errors could occur in both left and right directions (**Fig. 1a**). Therefore, depending on the CS-ON direction of individual PCs, the inward and outward errors will elicit CSs with high probability in those PCs whose CS-ON directions are aligned with the error vector (**Fig. 5a**, red circles), as compared to those cases in which the CS-ON direction and the error vector do not match^13,15,24,32,33^ (**Fig. 5a**, gray circles). In other words, for any retinal error in a particular trial, there will always be a subpopulation of PCs whose CS-ON direction matches the error vector, leading to CS-trials, and in others not, leading to ‘No-CS’ trials. For a given error in the n^th^ trial, we looked at its influence on the entire population of PCs in our data set and the consequences for the SS manifolds of the n+1^th^ trials, rather than restricting our analysis to only CS-ON units (see **Supplementary fig. 7a**), assuming that the behavior is based on the concerted action of both subpopulations. To this end, we combined trials following CS-trials from the pool of CS-ON PCs (i.e., Post-CS trials) and ‘No-CS’ trials from CS-OFF PCs (‘Post-No CS trials’), separately for outward (**Fig. 5b, left**) and inward errors (**Fig. 5b**, **right**). Importantly, we included all ‘CS-trials’ from CS-ON PCs (regardless of whether the actual error occurred or not), assuming that every CS in the error time window of 50-175 ms after the saccade was fired to report an error (referred to as simulated error trials in **Fig. 5**).

We found that CS firing associated with inward and outward errors modified the resulting PC-SS manifolds, based on PV as the kinematic-dependent parameter, differently (**Fig. 5c, top**). Relative to the ‘Post-No-CS’ trials, the normalized slope angle, capturing changes in PV-dependent manifold size relative to the rotation speed, profoundly increased in the post-inward error trials but decreased, albeit only slightly, for post-outward error trials (**Fig. 5c**, **bottom**). Could it be that this result may be influenced by the actual error-direction, rather than error-type? Our analysis comparing inward and outward errors made in the same direction revealed that the PC-SS manifolds of subsequent trials maintained their specificity for inward and outward errors, even if their vectors pointed in the same direction (**Supplementary fig. 7b-d**). Given that the PC manifold size and speed of the latent dynamics encode PV and saccade duration almost independently (**Fig. 4p,q**), this result suggested that CSs associated with inward and outward errors, potentially engaging the same population of PCs, tuned the population firing more towards duration coding in post-inward error trials (more compensatory duration change given PV) and PV coding in post-outward error trials (more compensatory PV change given duration).

Therefore, one would expect to see a reduction in subsequent saccade’s duration if a CS signal reporting an inward error, caused by an overshooting saccade, in the previous trial was present. On the other hand, in case of an outward error (i.e., undershooting saccade), CSs should trigger an increase in the PV of the next trial to reduce endpoint error. Indeed, this is what we found. When comparing the movement velocity of saccades accompanied by a CS to post-CS saccades, we observed that outward errors (undershooting) were corrected mainly through increasing the PV of the subsequent saccade with a slight increase in the velocity at the end of the saccade (**Fig. 5d**, **left**). In contrast, inward error-encoding-CSs prompted a significant decrease in the duration of the subsequent saccade, reflected by the narrowing of its velocity profile (**Fig. 5d**, **right**).

### Linear feed-forward network model shows high-dimensional transformations by the cerebellar cortex

We demonstrated that, despite the similar limit-cycle-like properties of MF and PC manifolds, they also exhibited crucial differences in their encoding of kinematic parameters. The climbing fiber-driven CSs clearly explain some of the differences between the two (**Fig. 5**, **Supplementary fig. 6**). However, additional inputs to PCs arriving from interneurons may also play a significant role.

Yet, we found that a linear feed-forward network (LFFN) from MFs to PCs^34^ (**Fig. 6a**) predicted the kinematics-independent and dependent activity components of all individual PCs with high fidelity (*R^2^*=0.984±0.018, mean±SD) (**Fig. 6b,c**), which allowed us to successfully reproduce the PC-SS manifolds from the MF activity (**Fig. 6d**; see also **Supplementary fig. 8a,b,c**). But how is it possible that already a simple linear transformation can explain the many differences between MF and PC-SS manifolds? This paradox led us to examine how many dimensions of MF (*d_MF_*) firing are necessary to make good predictions of the PC manifold.

**Figure 6.**
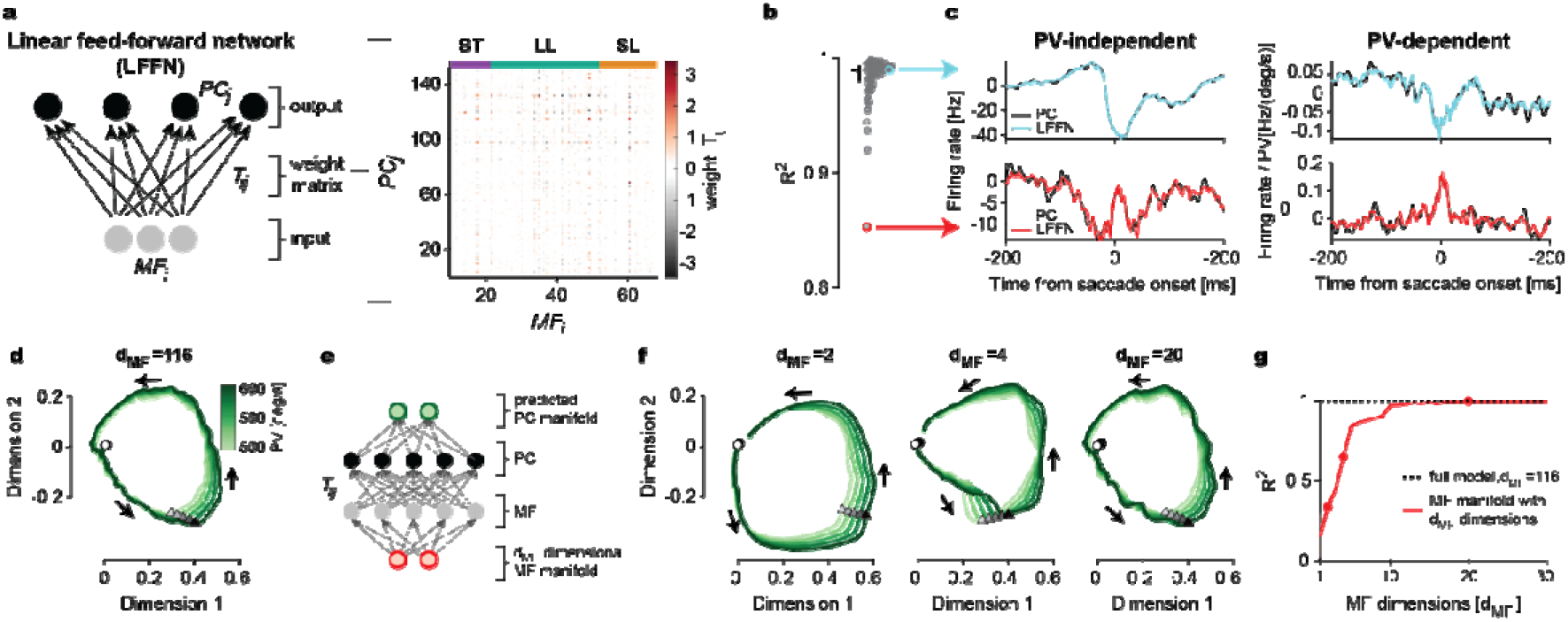
Linear feed-forward network (LFFN) model from MFs to PCs. **a** Left: Schematic diagram showing LFFN for MF-to-PC firing rate transformation. Right: weight matrix computed from the data. **b** Goodness of fit (R^2^) for individual PCs. The horizontal and vertical bar represents the median and from the first to third quantile, respectively. Colored circles correspond to examples shown in c. **c** Firing rates of example PCs (black, Top and Bottom) and LFFN predictions (color). The baselines are subtracted in the PV-independent component (Left). **d** LFFN prediction of PC manifolds in Fig. 4d. **e** Schematic diagram of the LFFN model for MF manifold-to-PC manifold transformation. **f** Examples of the predicted PC manifold from e when MF manifold dimension is *d*_MF_=2 (Left), 4 (Middle), and 20 (Right). **g** Goodness of fit for the predicted PC manifold to the data versus the input MF manifold dimensions *d*_MF_. Dots represent examples in f. Data are mean±SEM.

We addressed this question by two approaches, both leading to the conclusion that the number of dimensions that need to be considered while trying to account for the properties of MF activity is definitely much smaller than the maximum number of dimensions, *d_MF_*=116 (corresponding to the number of MFs in our data), but significantly higher than two or four, the dimensionalities capturing a major chunk of cell-to-cell variability in MFs and PCs, respectively (**Supplementary fig. 5a-h**). In the first approach, we first created the *d_MF_*-dimensional pseudo-population firing of MFs (**Fig. 6e**, grey circles) using the *d_MF_*-dimensional MF manifold (red circles), then generated the prediction of individual PC-SS firings using the LFFN (black circles), and finally identified the predicted PC-SS manifold (green circles). The PC-SS manifold (in the first two dimensions) was relatively poorly predicted (*R*^2^<0.9) when *d*_MF_<9 (**Fig. 6f,g**). In the second approach (**Supplementary fig. 8d**), we directly tested whether the prediction of individual PC firings requires high dimensional components in MF firings by another LFFN model, where MFs and PCs communicate through a dimensionally reduced submanifold, called the ‘communication subspace’^35^. This model also showed that a good prediction of individual PC responses requires a high-dimensional (*d*>15) communication subspace (**Supplementary fig. 8d,e**). Note that, the dimensions higher than four (i.e., *d*>4) explain only 4.3% of the total MF-to-MF variance together due to rapid decay (∝1/*d*^3.23^) in the explained variance (**Supplementary fig. 5b**). Therefore, the properties of PC-SS manifolds emerge as a consequence of a transformation by the cerebellar cortical circuit that amplifies those small variabilities in MF inputs.

## Discussion

The present study demonstrated the presence of multi-dimensional manifolds, latent in the activities of the cerebellar input and output, MFs and PCs respectively, and how their geometric and dynamic features encode key kinematic eye movement parameters. Climbing fiber-driven CSs, signaling error-related information to PCs, modify the PC manifolds, differentially depending not only on the direction of error but also the type of error, which predicts how the subsequent eye movements are corrected. Finally, we showed that the cerebellar cortical circuit amplifies seemingly insignificant variabilities in the MF activity to generate highly selective PC outputs.

The fast and repetitive nature of our paradigm induced cognitive fatigue, a gradual decline in the speed of saccades, which was compensated by duration upregulation^8^. However, on top of fatigue, we also observed natural trial-to-trial changes in the saccade velocity requiring rapid duration adjustments in order to guarantee endpoint precision. Therefore, the same velocity-duration trade-off mechanism that maintained movement accuracy across hundreds of trials within a session also ensured reduced endpoint variability (motor noise) on a trial-to-trial basis. The residual motor noise led to tiny, albeit specific error types, directed inward and outward respectively, depending on whether the eye movements were too large or fell short relative to the target location.

Depending on the firing pattern of individual MFs and SSs of PC units, we could broadly classify them into different categories by using strict statistical criteria to compute population averages of each category^13,26,36^. Yet, in our analysis, these units appeared continuous in their distribution (**Supplementary fig. 2**) rather than forming discrete clusters, due to a large cell-to-cell variability exceeding between-category distances. Therefore, one may question the reliability of the classify-and-average approach in testing the encoding of specific kinematic parameters as it may be prone to the risk of sampling bias. This problem gets even worse if one additionally considers the large between-session variability in eye movements also influencing the firing rates of individual units. To avoid exactly these biases, we estimated the firing rates of all individual units, based on a firing rate model that varies linearly with key kinematic parameters, to obtain a “pseudo-population” of MFs and PC-SSs. This allowed us to identify multi-dimensional, limit cycle-like manifolds of neuronal activity from these pseudo-populations capturing a significant proportion of cell-to-cell variability^22^.

PC discharge, the output of the cerebellar cortex, is only a few synapses away from the final stage motor neurons. Therefore, moving up the cerebellar circuitry, one would expect the PC signals to be far more refined and informative about the movements than the signals at earlier stages, e.g., at the level of MF afferents. At first glance, our results from the population analysis seemed to contradict this expectation as the MF pseudo-population exhibits a much more precise encoding of relevant kinematic parameters while PC-SS pseudo-population responses are sloppy and contaminated by a large heterogeneity in their firing patterns. However, a very different perspective is opened if one resorts to the low-dimensional pseudo-population manifolds that reveal the hidden dynamics of PC-SS activity for the flexible control of key movement parameters like velocity and duration in a movement phase specific manner. Furthermore, the PC manifolds carried significantly more disentangled representations of movements than the MF manifolds. Unlike MFs, the PC-SS manifolds exhibited distinct geometric and dynamical properties related to the two specific kinematic parameters, velocity and duration. This conspicuous difference between the MF and the PC-SS manifolds indicates a highly nontrivial transformation by the network.

Where do these differences stem from? Notably, our simple model, LFFN, simulating the MF-to-PC pathway could accurately explain the MF-to-PC transformation at the firing rate and manifold level, but only if the high dimensional components in the MF inputs, representing a tiny fraction (<5%) of the total MF-to-MF variability, were preserved. This result suggests that a disentangled movement encoding at the PC level emerges through substantial amplification of those seemingly insignificant variability of MF responses by the cerebellar network. Highly correlated activity, resulting in an apparently small dimensionality, has been widely observed in work on the cerebellar input layer^37–39^ (but see also ref.^40^). We found the same in our MF data, but our analysis together with the PC data suggests that enhancing small input variabilities is a fundamental information processing property of the cerebellar network. Furthermore, together with the finding that serial single-unit recordings are sufficient to generate reliable MF and PC manifolds, the prediction power of the LFFN model implies that MFs should use asynchronous firing rate coding.

PCs are also influenced by the direct climbing fiber pathway, imparting plastic changes in their activity via CSs. Indeed, we found that CSs modulated the geometry and dynamics of the PC-SS manifolds, on a trial-to-trial basis, in an error-type dependent manner, predicting selective post-CS parametric adjustments of eye movements. The forced error-based short-term saccadic adaptation is similarly error-type dependent^6^, which supports that PCs, by duration coding, control movements flexibly in response to external and internal (fatigue) changes^8^. On the other hand, recent studies have demonstrated the effects of CS-driven plasticity on the movement velocity, thereby emphasizing velocity-coding by PCs^13,15^. We demonstrated that those two mechanisms coexist and can be interwoven to exhibit complex forms of population-level plasticity. Therefore, the multidimensional nature of cerebellar computations is necessary for the flexible, context-dependent control of movements and their rapid adaptation.

The success of the linear model in describing the amplification of the variance as a consequence of the transformation of the MF input—not considering climbing fiber activity— indicates that the amplification of variance is independent of input from the inferior olive. However, this amplification is undoubtedly the basis allowing the climbing fiber system to select those chunks of information needed to optimize the movement.

## Materials and methods

### Animals, preparation, and surgical procedures

Two healthy male rhesus macaques (*Macaca mulatta; monkey K and monkey E, age: 10 years and 8 years, respectively*), purchased from the German Primate Center in Göttingen, were used for the purpose of this study. All data presented in this study were collected from these two animals using procedures that strictly adhered to the rules defined by the German as well as the European law and guidelines that were approved by the local authority (Regierungspräsidium Tübingen, veterinary license N7/18 and N4/14) and National Institutes of Health’s *Guide for the Care and Use of Laboratory Animals*. All training, experimental and surgical procedures were supervised by the veterinary service of Tübingen University.

As a first step, the animals were subjected to chair training which began in the animal facility where animals were encouraged to voluntarily enter a customized mobile chair for the first few weeks following which they were transported to the experimental area where they were gradually acclimatized to the new environment. To proceed with experimental training, it was necessary to painlessly immobilize the head in order to record eye movements reliably. Therefore, once the animals felt fully comfortable in the experimental setups, the first major surgical procedure of installing the foundations of cranial implants was performed. During this procedure, the scalp was cut open and these foundations, made out of titanium, were fixed to the skull using titanium bone screws. The scalp was then closed with the help of sutures under which the foundations were allowed to rest and stabilize for a minimum period of 3-4 months to ensure their durability and also full recovery of the animals. After this period, the second surgical procedure was performed in which the scalp was opened just enough to allow a titanium-based hexagonal tube-shaped head post to be attached to the base of the implanted head holder. Since this procedure was rather quick, the surgery was also accompanied by implantation of magnetic scleral search coils^41,42^ to record high-precision eye movements. After 2-3 weeks of recovery, monkeys were trained further on the behavioral task until their performance was accurate enough to consider neural recordings. To this end, the final surgical procedure was performed in which the upper part of the cylindrical titanium recording chamber (tilting backward by an angle of 30° with respect to the frontal plane, right above the midline of the cerebellum) was attached to the already implanted chamber foundation. A small area of the skull within the confines of the chamber was removed to allow electrode access to our region of interest, the oculomotor vermis (OMV, lobules VIC/VIIA). The position and orientation of the chamber were carefully planned and confirmed based on pre-and post-surgical MRI, respectively. All surgical procedures were performed under aseptic conditions using general anesthesia in which all vital physiological parameters (blood pressure, body temperature, heart rate, pO_2_ and pCO_2_) were closely monitored^43^. After surgery, analgesics (buprenorphine) were delivered to ensure painless recovery which was monitored using regular ethograms under the strict supervision of animal caretakers and veterinarians.

### Experimental setup and behavioral task

All experiments were performed inside a dark room where monkeys, with their heads fixed, were seated comfortably in a primate chair placed at a distance of 38 cm in front of a CRT monitor such that their body axis was aligned to the center of the monitor. All neural and behavioral data presented in this study were collected during a simple to-and-fro saccade task in which monkeys were asked to rapidly shift their eye gaze repeatedly in order to follow a jumping target that appeared in two fixed locations along the horizontal axis on the monitor in an alternating manner (**Fig. 1a**). Before the beginning of each trial, the fixation target (a red dot of diameter 0.2 deg) appeared at the center of the monitor with an invisible fixation window of size 2×2 deg centered on it. Only if the monkeys moved their gaze within the fixation window the trial was initiated. This was followed by a short fixation period ranging from 400 to 600 ms from trial onset after which the fixation target vanished and, at the same time, another target (with the same properties as the fixation target) appeared at a new horizontal location, giving the impression that the target “jumped” centrifugally (**Fig. 1a**, solid arrows), i.e., from the center of the screen to this new location. The size (=15 deg) and the direction (left or right) of the target jump were kept constant within a session. Every target jump served as a ‘go-cue’ which prompted the monkey to execute a saccade towards the new target location within the 2×2 deg fixation window centered on it, in order to receive an instantaneous reward (water drops) marking the end of a trial. The peripheral target disappeared approximately 700-900 ms relative to the go-cue, immediately after which the central fixation dot reappeared indicating the beginning of the next centrifugal trial. In order to proceed with the next trial, the monkey made a saccade from the peripheral target back to the central location (i.e., centripetal saccade, see dashed arrows in **Fig. 1a**). In other words, the appearance of the central fixation dot served as a go-cue for centripetal saccades, although these saccades were not rewarded. Depending on the motivation of the monkeys to perform the task, as well as the duration for which a PC could be kept well isolated, the number of trials varied in each session (median=307 trials) with each trial lasting for 1200 ms. While the fatigue-inducing fast and repetitive nature of the paradigm allowed us to capture both trial-by-trial and gradually declining changes in the peak velocity of centrifugal and centripetal saccades, the natural endpoint variability in saccades, on the other hand, observed as over-or undershoots resulting in inward (**Fig. 1a**, see yellow arrows) or outward errors (**Fig. 1a**, see green arrows), allowed us to measure the CS’s preferred and anti-preferred direction of error for an individual PC. All experimental parameters were designed and controlled using in-house Linux-based software, NREC (http://nrec.neurologie.uni-tuebingen.de).

### Electrophysiological recordings, identification of Purkinje cells and mossy fibers in the oculomotor vermis

All electrophysiological recordings of PCs (n=151) and mossy fibers (n=117) from the OMV were performed using glass-coated tungsten microelectrodes (impedance: 1-2 MΩ), manufactured by Alpha Omega Engineering, Nazareth, Israel. To target the OMV, as predicted by the MRI scans, the position of electrodes along the rostrocaudal (i.e, Y-axis) and lateral (i.e, X-axis) axis were manually adjusted with the help of a custom-made microdrive, temporarily mounted on the recording chamber during each experimental session. The depth of the electrode was controlled using a modular multi-electrode manipulator (Electrode Positioning System and Multi-Channel Processor, Alpha Omega Engineering). The exact location of the OMV was confirmed based on careful inspection of online audio-visual feedback of the electrode signals, reflecting multi-unit granule cells activity, that exhibited strong modulations in response to fast eye movements.

For PC recordings, extracellular potentials sampled at 25 KHz were high (300 Hz- 3 KHz) and low (30 Hz-400 Hz) band-pass filtered to obtain action potentials and LFP signals, respectively. Individual PC units were identified based on the presence of two types of action potential signals, high-frequency simple spikes (SSs) and low-frequency complex spikes (CSs), the latter characterized by a polyphasic wave morphology in the action potential trace paralleled by large deflections in the LFP signals. The fact that both signals originate from the same unit was confirmed online by the suppression of SS discharge for 10-20 ms when aligned to the occurrence of a CS^44–46^. Although the final characterization of CSs was based on an offline neural networks approach^47^, we relied on the performance of Alpha Omega Engineering’s Multi Spikes Detector for detecting SSs online.

In order to record from mossy fibers (MF) in the granular layer, we adjusted the upper cut-off frequency of the high band-pass filtered to 5 KHz while keeping the lower cut-off frequency the same as 300 Hz. The identification of MFs was based on their strong directionally selective response to saccades, firing up to several hundred spikes per second in the preferred direction and seldomly in the opposite direction. Unlike the relatively longer duration SSs (mean duration: 1.5 ms), MF units exhibited much shorter duration (mean duration: 0.6 ms), mostly mono- and biphasic shaped waveforms while occasionally exhibiting a negative after-wave^16,25,26,48,49^. Additionally, MFs exhibited a wide range of inter-spike intervals^16^ (mean ± sd: 82.7 ± 86 ms) as compared to those of PC SSs (mean ± sd: 19.5 ± 2.6 ms).

### Classification of mossy fiber responses

Unlike the bidirectional SS discharge of PCs, well-isolated MF units exhibited a strong and clear preference for saccades made in one of the two horizontal directions. This property allowed us to pre-determine the preferred direction of the MF unit under investigation and use that direction as the rewarded direction in which the centrifugal saccades were made. A majority (115 out of 117) MF units investigated in this study exhibited a much stronger (“burst-type”) discharge during the peri-saccadic period in their preferred direction (=centrifugal direction) as compared to the opposite, non-preferred direction (=centripetal direction) in which very few or almost no spikes fired, resulting in weak modulations. Therefore, MF responses only in the centrifugal direction were considered for classification and all analyses. In the other 2 units, we did not observe a peri-saccadic burst

Overall, we observed two main types of burst modulations: the eye position-related tonic discharge preceded by a saccade-related burst, i.e., the ‘burst-tonic’ type, and the saccade-related burst discharges that remained mostly silent outside the peri-saccadic period, i.e., ‘phasic’ type. In order to identify the ‘burst-tonic’ responses, we first identified those units in which the difference between the average firing rate in the post-saccadic period (150 to 250 ms from saccade onset) and the pre-saccadic period (−250 to −150 ms from saccade onset) was larger than 1.5 x standard deviation (SD) of the average firing rate during the pre-saccadic period. Next, we compared the slope values of the linear regression fits applied on the pre-and post-saccadic firing responses, and only those cases in which no significant difference between the slopes was observed, were labeled as ‘burst-tonic’ responses (n=24; **Fig. 2a,b**; see BT). In other words, if the post-saccadic MF activity was not only larger than the pre-saccadic activity but also remained elevated after the saccade-related burst discharge, the unit’s response was classified as a ‘burst-tonic’ type. The ‘phasic bursts’, on the other hand, were further categorized into ‘long-lead burst’ types and ‘short-lead burst’ types, based on the timing of each MF unit’s burst modulation onset relative to saccade onset^26^. For this, modulation onsets were detected whenever the averaged MF response crossed a threshold (defined as 3 x SD of baseline activity during −400 to −200 ms from saccade onset). To this end, all MF units in which the burst modulation led the saccade onset by more than 15 ms were labeled as ‘long-lead burst’ types (n=60; **Fig. 2a,b**; see LLB), whereas those that started firing less than 15 ms before the saccade onset were classified as ‘short-lead burst’ (SLB) types (n=27; **Fig. 2a,b**; see SLB). The value 15 ms was chosen, based on the observed SD value of modulation onsets of ‘long-lead burst’ MF units identified by Ohstuka and Noda^26^. Given that the timing of the detected modulation onsets was a crucial factor in separating these two categories, in addition to the clarity of their firing patterns, spike data were not smoothened using a Gaussian kernel, as in the case of SSs. Based on this criteria, 4 units (in addition to 2 non-bursting units) could not be categorized into any of the three categories as in those cases the onset of burst modulation occurred after (i.e., lagged) saccade onset.

### Classification of simple spike responses

SS responses of individual PCs were broadly categorized into 4 types—burst, pause, burst-pause and pause-burst—based on their pattern of firing during the perisaccadic period of −50 to 150 ms from the end of primary saccades (note: all primary saccades between 13 to 17 degrees of amplitude were detected using a velocity threshold of 30 deg/s). To this end, we estimated the mean spike density function of the SS discharge of individual PCs by first convolving the time of each SS event detected within a trial with a normalized Gaussian kernel (sd=5 ms) and then averaging across all trials.

Given that centrifugal and centripetal saccades were made horizontally in opposite directions, their SS firing patterns could be entirely different. For instance, a PC could demonstrate a sharp peri-saccadic increase (or burst) in SS firing for a rightward centripetal saccade, whereas in the opposite direction (i.e., left centrifugal) the same PC could exhibit a sudden drop in SS firing (or pause). Therefore, each PC’s SS response was characterized by two response profiles (one for each tested direction) and both were considered independently, as separate units (n=302; 151 PCs x 2 directions), in our classification procedure described below.

As the first step, we used threshold-based criteria to label each SS response with one of the four types based on the polarity of the response modulation. For this, we identified all maximum (peaks) and minimum (troughs) SS firing rates (detected using the MATLAB function ‘findpeaks’, minimum peak distance = 10 ms, minimum peak prominence = 2 spikes/s) during the peri-saccadic period. The modulation was considered significant if the peaks and troughs crossed an upper and a lower threshold (defined as ± 5 x s.d of baseline activity during the −250 to −100 ms from saccade onset), respectively. The SS response was classified as a ‘burst’ or a ‘pause’ type if we encountered only a monophasic increase or decrease in SS firing during the peri-saccadic period. Responses were categorized into ‘burst-pause’ or a ‘pause-burst’ types if the first modulation in the biphasic responses showed an increase (followed by decrease) or a decrease (followed by increase) in SS firing, respectively. Next, we ran a principal component analysis (PCA) on the 302 SS responses (CF and CP combined) to obtain a 2D plot (**Fig. 3a**) of their first two principal components (explaining 62.2% of the total variance) that seemed to appear as overlapping clusters organized in a circular pattern, centered around the origin. For better discrimination of these clusters, we relied on the SS response labels (identified in the first step) to obtain decision boundaries by resorting to linear discriminant analysis (LDA). As shown in **Fig. 3a**(dashed lines), the first decision boundary separated the ‘burst’ (blue cluster) from ‘burst-pause’ (green cluster) types, as well as the ‘pause’ (orange cluster) from the ‘pause-burst’ (red cluster) types. On the other hand, the second decision boundary separated the ‘burst’ from ‘pause-burst’ types, and also the ‘pause’ from the ‘burst-pause’ types. As compared to the threshold-based labeling of these response patterns, the LDA approach was clearly better in separating these response types (**Supplementary fig. 2c,d**).

### Rate models for individual MFs and PCs

We constructed the firing rate model of individual MFs and PCs by using a linear combination of kinematics-independent and kinematics-dependent components. Given the baseline-subtracted dynamic firing rate of the *i*^th^ unit, *R*_i_(*t*,**z**), where *t* is the time from saccade onset and ***z*** is a vector of the specific movement kinematic parameter (e.g., **z**= [PV] or [duration], or a pair of kinematic parameters, i.e., **z**= [PV, duration]), we modeled the firing rate vector of a “pseudo-population” containing *N* number of neurons, **R**(*t*,**z**) = [R_*1*_(*t*,z); R_*2*_(*t*,z);…; R_*N*_(*t*,z)], as

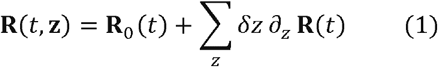

where **R**_0_ and ∂_*z*_**R** are the kinematics-independent and dependent part, respectively, and *δZ*=*z-z_0_* is the deviation of *z* from the mean value of *z, z_0_*. We used the multivariate linear regression of the firing rate data with respect to the kinematic parameters (**Supplementary fig. 3a**) for each unit to find the model components for all unit data (**Supplementary fig. 3b,c**). See **supplementary methods** for details.

### Estimation of manifolds

To find the dimensionally reduced approximation of the the population rate model, **R**(*t*,***z***) in Equation 1, given by **R**_0_ and ∂_*z*_**R**, we followed the following steps: First, **R**_0_ and ∂_*z*_**R** were converted to (*N*,*T*) matrices by discretizing time where *T* is a length in time in msec. We performed PCA on **R**_0_, which is the firing rates at ***z*** = ***z***_0_. We obtained a dimensionally reduced representation, a manifold, **P**_*K*_ such that **R**_0_ ≈ **WP**_*K*_ where **W** is some (*N*,*K*) dimensional matrix (*K<N*). We determined *K* by finding the number of dimensions capturing >85% of the total variability and confirmed it by the cross-validation analysis. Then, we estimated the linear approximation of how the kinematics-dependent component, ∂_z_**R**, would change the PCA result of the firing rates if ***z*** deviates from ***z***_0_. Our analytic estimation showed that it is enough to consider a change in **P**_*K*_ as,

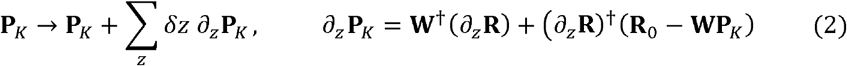

to predict the PCA results of **R**(*t*,**z**) with sufficient accuracy. ∂_*z*_**W** is a matrix obtained in the second step and describes how the higher dimensional (>*K*) components move into the *K*-dimensional subspace when kinematic parameters change and † represents conjugate transpose. See **supplementary methods** for details.

### Analysis of manifolds

Given a manifold of MFs or PCs given movement kinematics, we computed the manifold size and rotation speed in 2D (**Fig. 4–5**). We defined the manifold size by an enclosed area within the circular trajectory in 2D, which is computed by numerically integrating the areas of triangles defined by two neighboring data points and the origin (0,0). For the rotation speed, we first computed the phase of rotation *θ* at each data point (*x*,*y*) by *θ* = tan^−1^(Δ*y*/Δ*x*) where (Δ*x*, Δ*y*) = (*x-x*_0_, *y*–*y*_0_) and (*x*_0_, *y*_0_) is a reference point defined by [(maximum of *x* coordinate data)/2, 0]. Then, we estimated the time *T_3/4_* from the trial beginning *t* = −250 ms, where *θ* ≈ −180° by definition, to the point *θ* = 90° (rotation of 3/4 cycles), finally finding the average rotation speed by 270°/*T_3/4_*. We summarized how the manifold size and rotation speed vary with the kinematic parameters by computing the normalized slope angle in the manifold size and rotation speed plane (**Fig. 4p,q** and **5c**). To do so, we first normalized the manifold size and rotation speed data for all cases by the standard deviations of the control case, which was the correlated variation in **Fig. 4p,q** and post-no-CS case in **Fig 5c**. The slope angle was computed in each case in the normalized coordinates. We also performed the comparison/alignment analysis of multiple manifolds using the canonical correlation analysis^23,27^. See **supplementary methods** for details.

### Linear feed-forward network models

The LFFN models had movement kinematics-independent and dependent components for output variables (**Y**, ∂_**z**_**Y**) and input (**X**, ∂_**z**_**X**), such as PC and MF firing rates in Fig. 6a-b. We assumed that the movement variable ***z*** follows the Gaussian distribution and estimated the weight matrix **T** to minimize the least-square error,

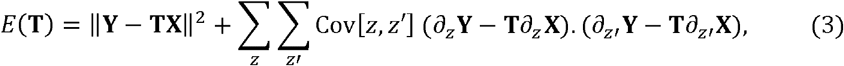

Performances of all the LFFN models were measured by this least-square error. To prevent overfitting we used the LASSO regression^50^ where the hyperparameter is chosen by AIC minimization. For the manifold-transformation LFFN (**Fig. 6e**), we reused **T** from the MF-to-PC LFFN model but replaced the input variables by those approximated by the *d*_MF_-dimensional MF manifold. The communication subspace model (**Supplementary fig. 8d,e**) was obtained by the rank-reduced regression^35^ with the error function in Equation 3. See **supplementary methods**for details.

### Statistical analysis

In most data analyses, we evaluated a mean and SEM by the jackknife resampling except for two quantities. In testing the prediction of the population-averaged firing rate by models (**Supplementary fig. 3e,g** and **4b,d**), we separated trials into two equal-sized sets, trained the model by only one of them (train data), and tested it on the other data set (test data). In **Fig. 6g**, we used the bootstrap procedure that randomly sampled the goodness of fit for individual time points and computed their averages with 500 repetitions to give the bootstrap mean and SEM.

---

**Supplementary figure 1.**
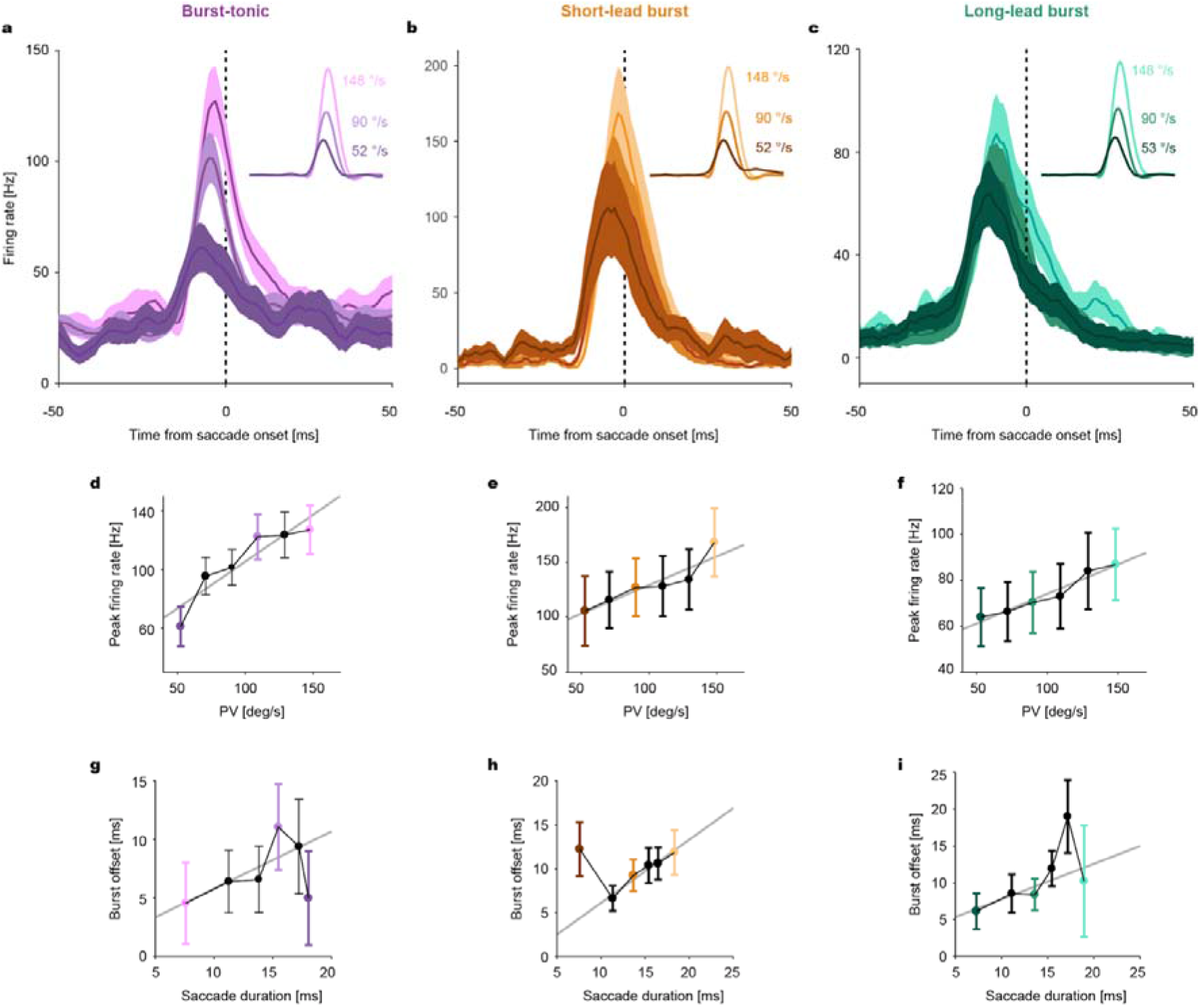
Linear encoding of microsaccades’ kinematics by mossy fibers (MFs). **a, b, c** Population response of burst-tonic (BT, purple), short-lead burst (SL, yellow) and long-lead burst (LL, green) MFs to saccades of different peak velocities (PV, see insets for average velocity profiles), represented by different shades. **d, e, f** Average peak firing rate as a function of saccade peak velocity (bin size=20 deg/s) for each MF category. Burst-tonic: p=0.012, R^2^=0.85; Short-lead burst: p=0.01, R^2^=0.85; Long-lead burst: p=0.001, R^2^=0.95. **g, h, i** Average burst offset relative to saccade onset as a function of saccade duration (calculated from velocity bins) for each MF category. Burst-tonic: p=0.18, R^2^=0.21; Short-lead burst: p=0.01, R^2^=0.02; Long-lead burst: p=0.22, R^2^=0.44. Solid gray lines represent the linear regression fits. Dark and light-colored bins correspond to the high and low peak velocity bins, respectively, for which population responses in a, b and c are plotted for comparison. Data are mean±SEM.

**Supplementary figure 2.**
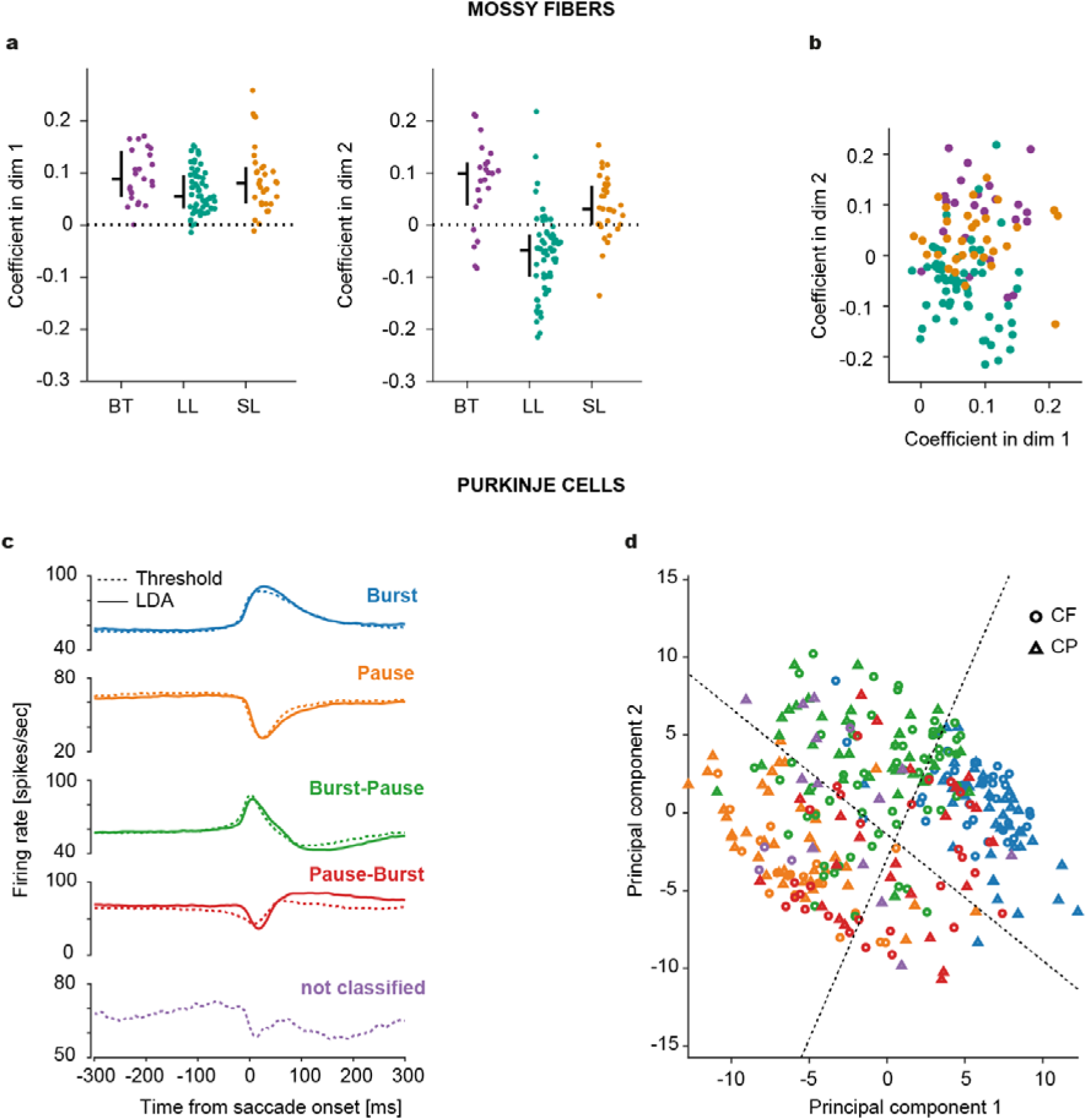
MF and PC units appear continuous in their distributions. **a** Coefficients of MFs for the first (Left) and second (Right) dimension in the MF manifold. Horizontal bar: median. Vertical bar: Range from the first to third quantile. **b** 2D scatterplot for the coefficients in a. Note a nearly continuous distribution of data points with significant overlaps between BT, SL and LL MF types (denoted by colors). **C** Average firing response of all PCs categorized into burst (blue), pause (orange), burst-pause(green) and pause-burst (red) types by threshold-based labeling (dashed lines) and linear discriminant analysis (LDA). Purple dashed lines indicate the average response of those PCs units which could not be classified into any of the four categories by the threshold-based method. **d** 2D scatterplot of the coefficients of first two principal components identified by the PCA for individual PC units recorded for centrifugal (CF, circles) and centripetal (CP, triangles) saccades. Dashed lines indicate the decision boundaries estimated by the LDA. Colors represent the PC category. Note, the overlap between different categories.

**Supplementary figure 3.**
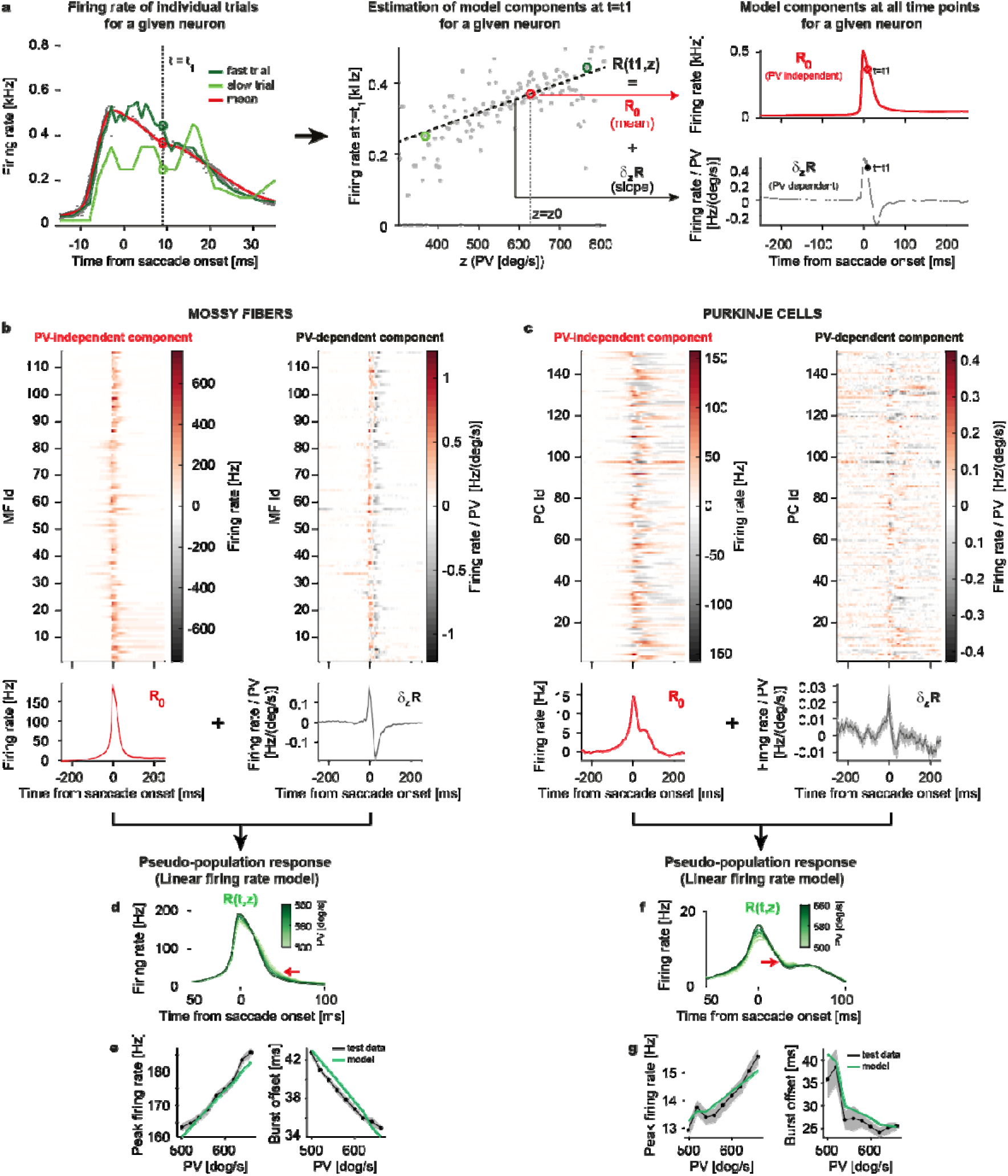
MF and PC-SS linear firing rate models. **a** Schematic illustration showing the steps involved in the construction of a rate model for individual (MF and PC) units/neurons using PV as the control kinematic parameter for the model. From the time-dependent firing rate estimations for individual trials of a given unit (Left), we create the linear regression model of movement kinematics, such as PV, versus firing rates at each time point (Middle). For example, given a linear dependence of MF or PC-SS firing rates on saccade PV, a randomly chosen saccade with high PV will be associated with higher firing rates (fast trial, dark green) as compared to a low PV saccade (slow trial, light green) and the difference between firing rates will be more pronounced during the initial phase of a saccade. In pre- and post-saccadic periods, where fast and slow trials can no longer be differentiated by PV, the differences in firing rates will also eventually disappear. There the slopes of regression will be much steeper at time points that fall within the peri-saccadic period. From the center (mean) and slope of the result, we obtain the kinematics-independent and dependent components (Right). **b,c** Top: Heat-map showing PV-independent (**R**_0_) and dependent components (∂_PV_**R**) for individual MF (b) and PC models (c). Bottom: Population averages. The baseline firing rates are subtracted. **d** Pseudo-population average firing rate for different PVs, computed from MF models in **b**. A red arrow indicates the point of burst offset. **e** Average peak firing rate (Left) and burst offset time (Right) vs PV from the models and test data. Goodness of fit: *R^2^* = 0.929±0.005 (Left), 0.887±0.026 (Right). **f,g** Same plots as d,e for PCs. *R^2^* = 0.809±0.023 (Left), 0.619±0.095 (Right). Note that using the PV-and-duration model did not significantly improve the predictions in e,g: peak firing rate vs PV, MFs: *R^2^*=0.929 ± 0.005, PCs: *R^2^*=0.791 ± 0.017; burst offset vs PV, MFs: *R^2^*=0.892 ± 0.021; PCs: *R^2^*=0.702 ± 0.05. Data are mean±SEM.

**Supplementary figure 4.**
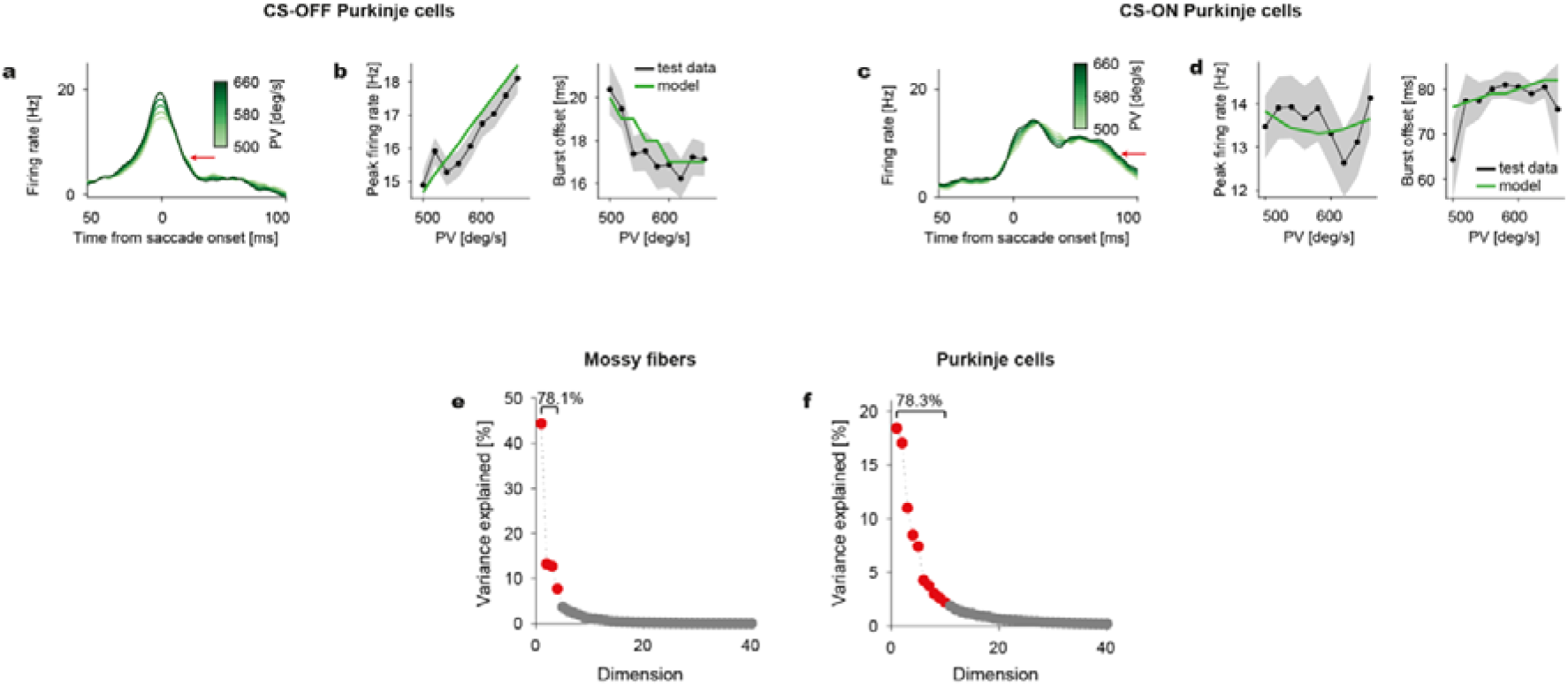
Pseudo-population SS response for CS-ON and CS-OFF population of PCs. **a** PV-dependent population average firing rates. The red arrow indicates the point of burst offset. **b** Average peak firing rate (Left) and burst offset time (Right) versus PV from the models and test data in CS-OFF direction. Goodness of fit: R^2^ =0.689±0.051 (Left), 0.433±0.121 (Right). **c** Same as a, but for CS-ON PCs. **d** The same plots as b for CS-ON PCs. R^2^ = 0.018±0.033 (Left), 0.092±0.088 (Right). The baseline rates are subtracted in all data. **e,f** Variance explained by each dimension in the PCA analysis of the PV-dependent components of the MF (e) and PC-SS models (f). Components with >78% are marked in red. Data are mean±SEM.

**Supplementary figure 5.**
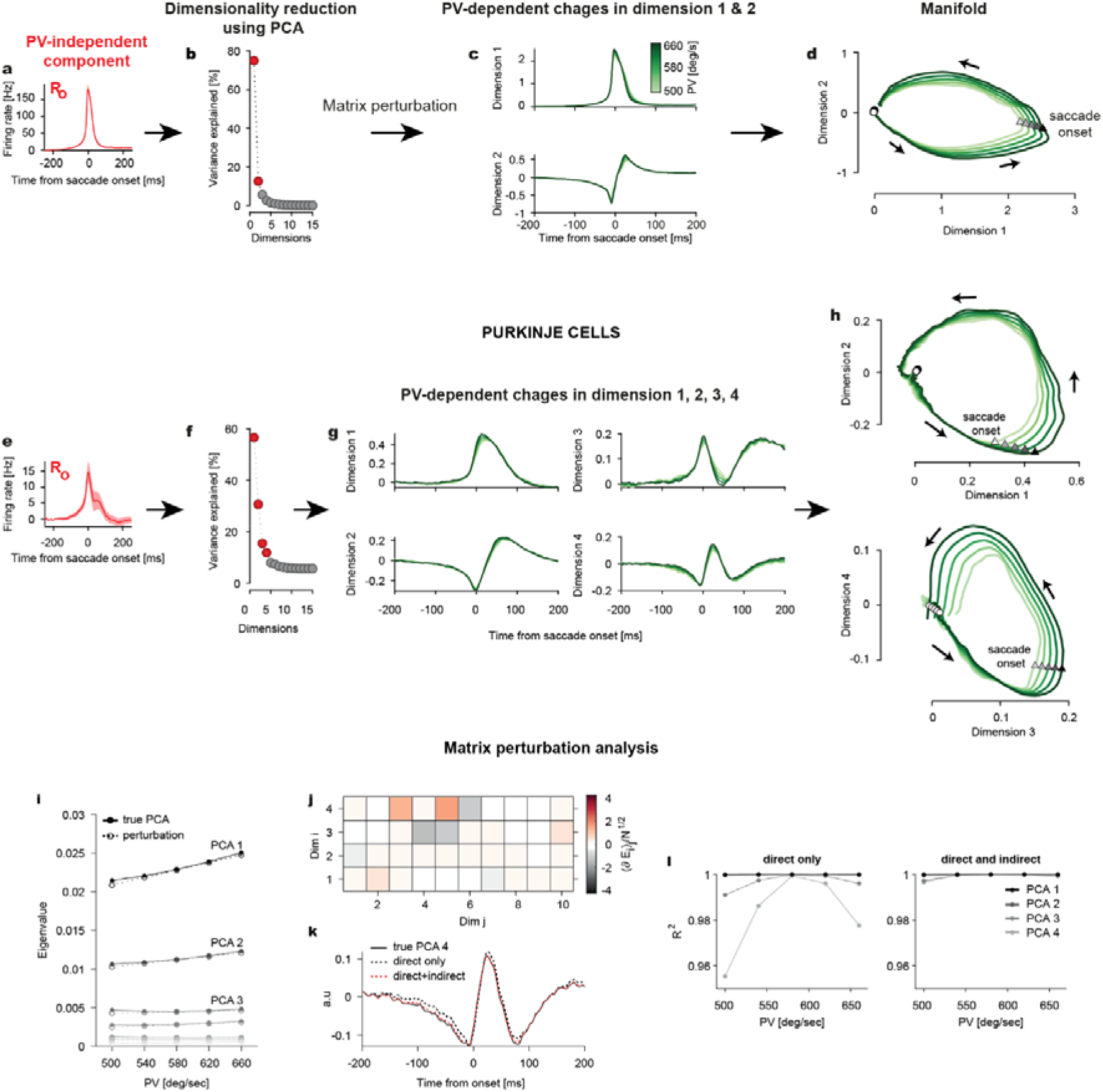
Step-by-step procedure for identifying manifolds. **a** Average of all the PV-independent components of all MF units (parameter: PV), which are subjected to PCA in the first step. **b** The first two principal components (or dimensions, red dots) of the PV-independent components explains a dominant fraction of cell-to-cell variability. **c** Matrix perturbation analysis (see Methods and Supplementary methods) computes PV-dependent changes in the first two dimensions, plotted against time. **d** Limit cycle-like 2D MF manifolds are identified by plotting the first two dimensions against each other for different values of PV (shades of green). Note how the manifolds increase in size, depicted by the separation of curves, with increments in PV, both before and after saccade onset (triangles). Arrows indicate the direction of rotation. **e-h** Same plots as a-d for PCs. Here, four dimensions explain a dominant fraction of cell-to-cell variability. Note how the differences in manifold, in the first two dimensions, are limited to periods after saccade onset, whereas in the third and fourth dimensions changes also appear before saccade onset. Note that the trajectories for the third and fourth dimensions (h, bottom) are plotted only until 50 ms after saccade onset to highlight the changes occurring before saccade onset. **i** Comparison of the eigenvalues from PCA (solid) of the PC-SS data and the prediction of the matrix perturbation theory (dotted). **j** Contribution of the original PCA eigenvectors (column) to their PV-dependent perturbative changes in each dimension (row), based on the PC-SS data. The coefficients (color) are normalized by *N*^1/2^ where *N*=151 is a number of PCs. **k** Fourth principal component of the PC-SS data when PV=500°/s (black solid), approximation only by the direct projection (dotted; see Supplementary Methods), approximation with an additional indirect projection from the eigenvector perturbation shown in j (red). **i** Accuracy of the approximated PCA components from the PC-SS data with different PVs, with only the direct projection (Left) and direct and indirect projection (Right).

**Supplementary figure 6.**
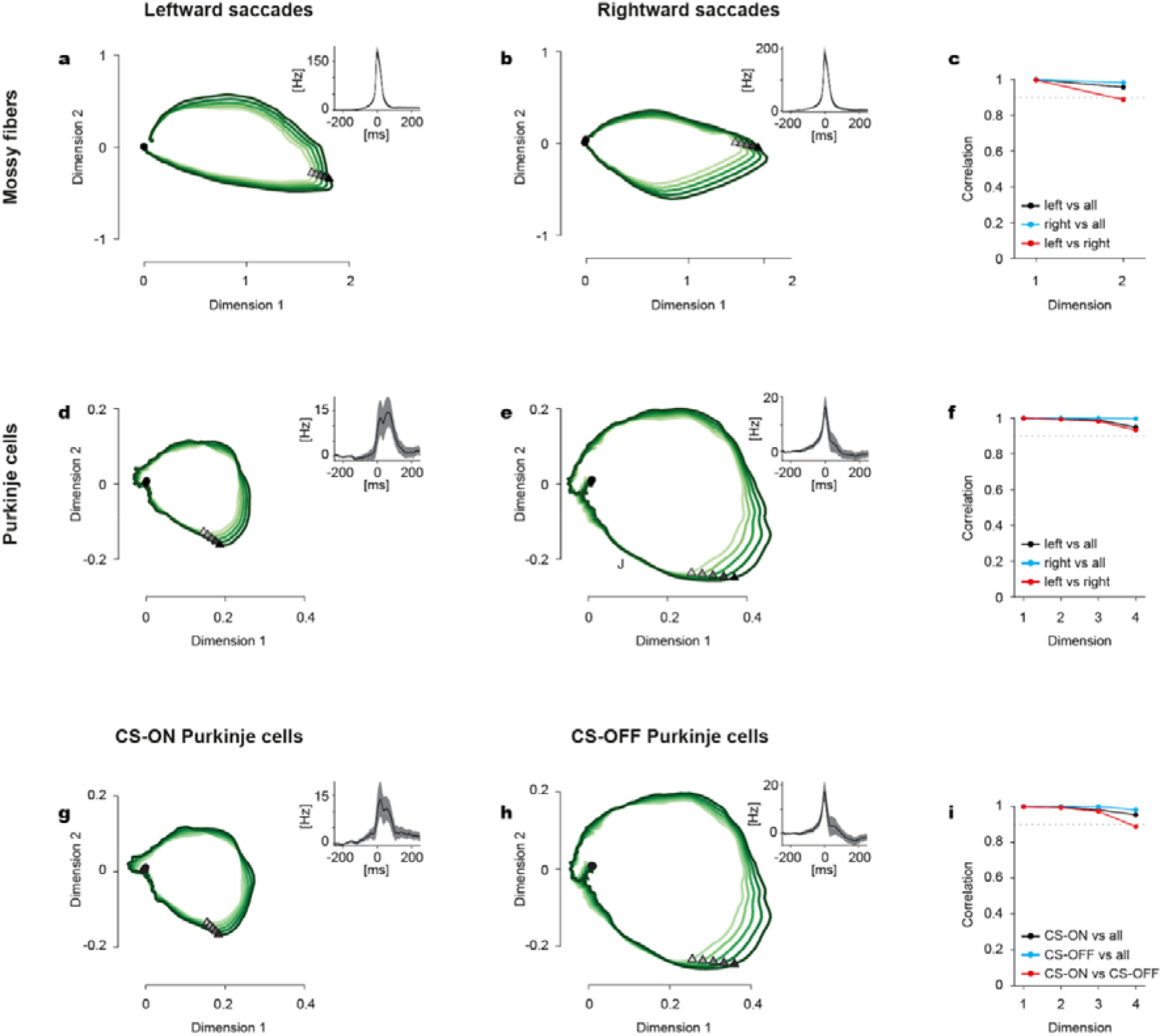
MF and PC-SS firing rate models and manifolds for different eye movement directions. **a,b** 2D plots of the MF manifolds from the left- and right-directed saccades. Insets show the population average of the PV-independent components of all MF firing rate models (control parameter: PV). **c** Canonical correlation of each dimension in the MF manifold between the left and right directions. Dotted line represents correlation=0.9. **d-f** Same plots as a-c for PC-SS manifolds. **g-h** 2D manifolds of PC-SSs separately for a population of CS-ON and CS-OFF PCs. Note that the similarity to d-f is due to that the CS-ON direction is mostly

**Supplementary figure 7.**
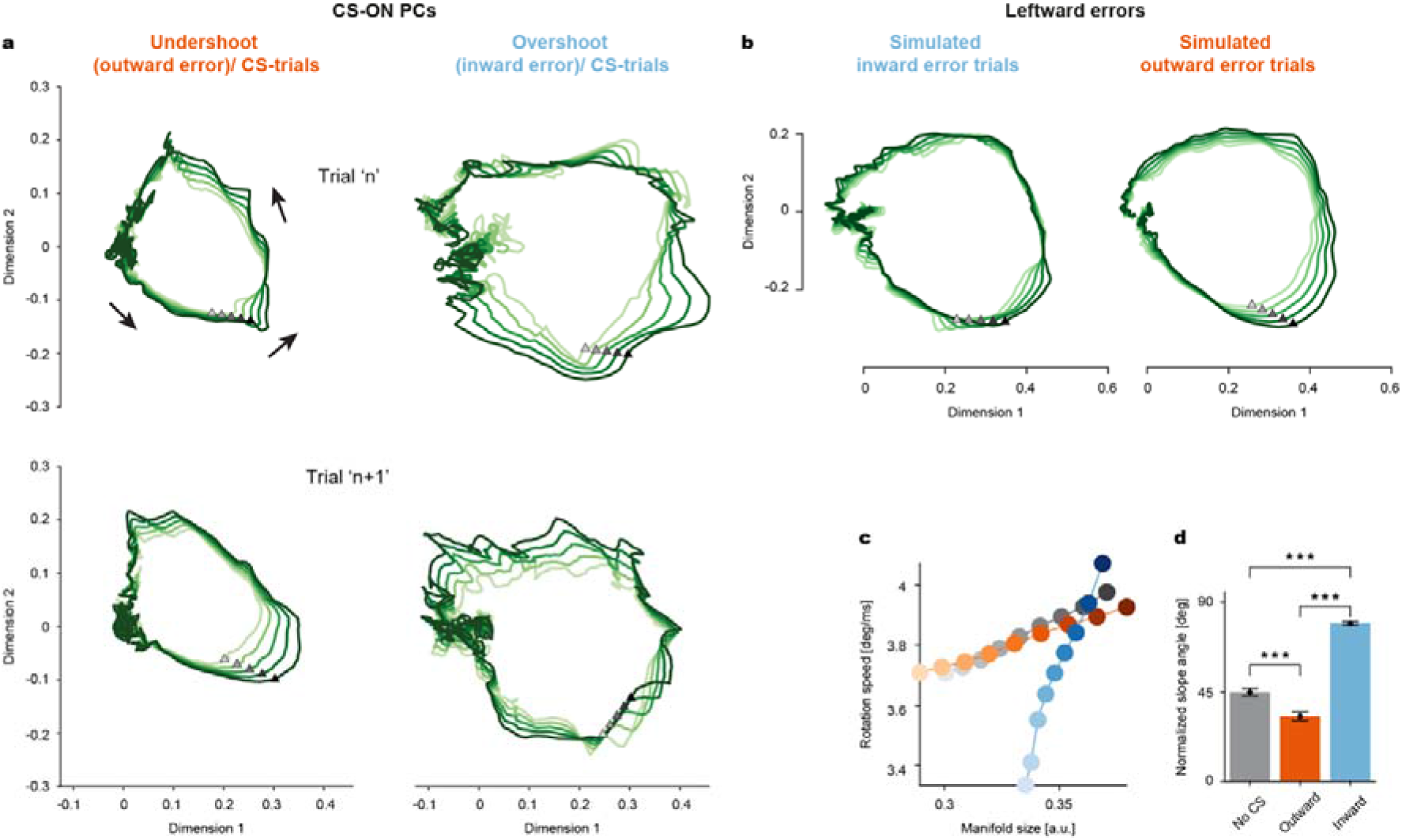
CSs influence the PC-SS manifolds differently depending on the type of error, even if the error direction is the same. **a** Up: Manifolds when outward (left) and inward (right) errors occurred in trial ‘n’ accompanied by CS firing in the post-saccadic period. Down: Manifolds in the subsequent trial ‘n+1’. Note how the manifolds in trial n+1 change differently for outward and inward errors, similar to those for simulated error trials shown in Fig. 5b. Filled triangles indicate the saccade onsets and the black arrows indicate the direction of rotation for all manifolds shown. **b** Manifolds for simulated post-inward and post-outward error trials controlled for error direction (i.e., Leftward errors). **c** Rotation speed as a function of manifold size for simulated post-inward (blue), post-outward (orange) and no-CS control (gray) trials. **d** A comparison of normalized slope angles for each condition. Note that the error-type specific changes in manifolds are preserved, i.e., an outward error-related increase in manifold size (indicated by the relatively flatter slope of the orange curve as compared to No-CS) and inward-error related change in rotation speed (indicated by relatively steeper slope as compared to the No-CS condition), despite the error vector pointing in the same left direction. T-value (No-CS, Outward) =5.93, p=9.88×10^−9^; T-value (Outward, Inward) = −28.28, p=1.59×10^−62^; T-value (No-CS, Inward) =−23.03, p=1.41×10^−51^.

**Supplementary figure 8.**
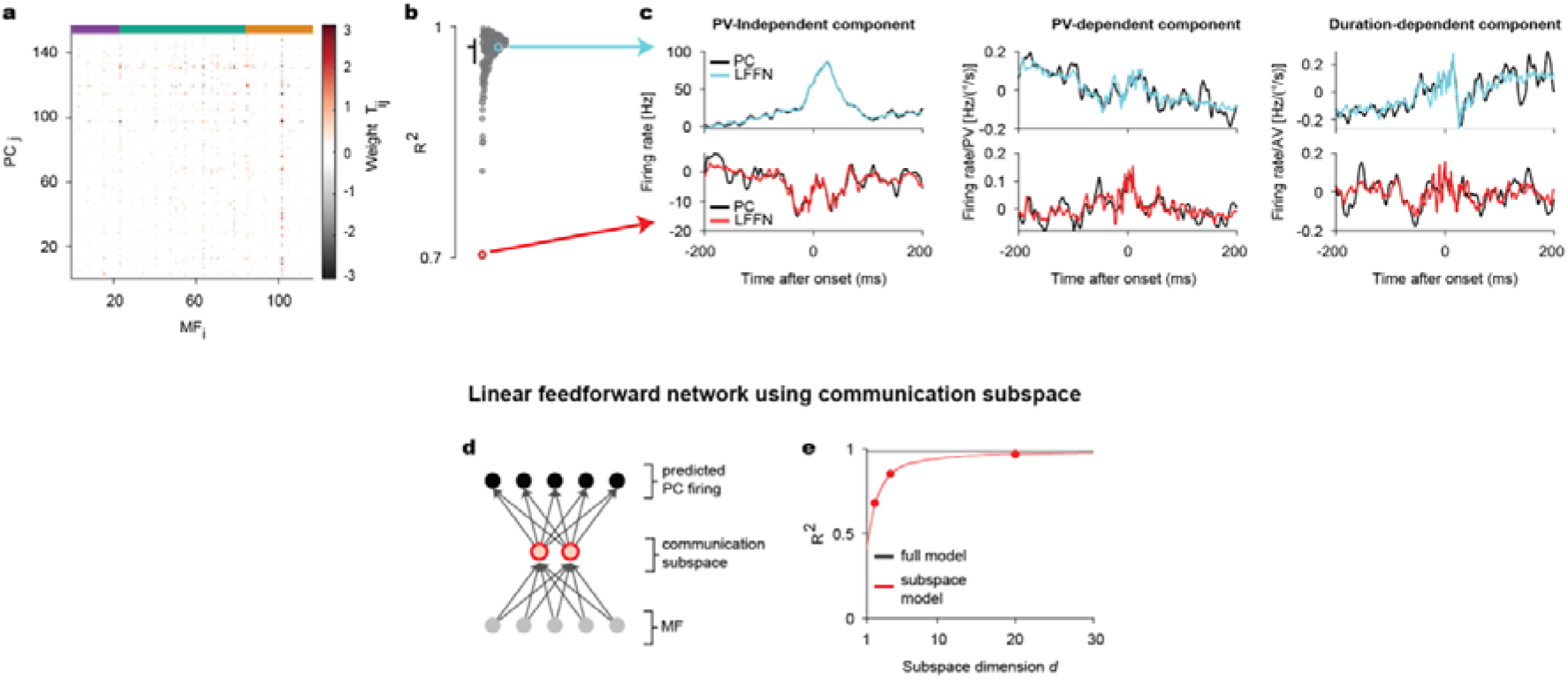
Linear feed-forward network (LFFN) model for MF-to-PC transformation with PV and duration dependence. **a** Weight matrix of the MF-to-PC network model. **b** Goodness of fit for individual PCs. Colored circles represent the examples in c. Horizontal bar: Median. Vertical bar: Range from the first to third quantile. **c** PV-independent (Left), PV- (Middle), and duration-dependent component of example PC-SS rate models (black) and prediction by LFFN (color). The baseline rates are subtracted in the PV-independent components. **d** A schematic illustration of the communication subspace model of MF-to-PC transformation. A communication subspace, of the dimensionality d, limits the feedforward network using all the dimensions in the MF rates to predict PC-SS rates. **e** Goodness of fit for model prediction of PC-SS rates. Red dots represent d=2, 4, and 20. Data are mean±SEM.

