## Supplementary information for "Multidimensional cerebellar computations for flexible kinematic control of movements"

### Supplementary Methods

#### Fitting the linear rate model to data

In the main text, we modeled the firing rate vector of a "pseudo-population" containing $N$ number of neurons, $\mathbf{R}(t, \mathbf{z}) = [R_1(t, \mathbf{z}); R_2(t, v); \ldots; R_N(t, \mathbf{z})]$, as

$$\mathbf{R}(t, \mathbf{z}) = \mathbf{R}_0(t) + \sum_z \delta z \, \mathbf{R}_z(t). \tag{S1}$$

where $\mathbf{R}_0$ and $\partial_z \mathbf{R}$ are the kinematics-independent and dependent part, respectively. $\delta z = z - z_0$ is the deviation of $z$ from the mean value of $z$, $z_0$.

We fitted this model to the MF and PC data in the following way: We first estimated the firing rate at each trial by the fractional interspike interval method[1] ($\tau$=5 ms). Other estimation methods, such as spike train smoothening by a Gaussian kernel, did not change the results. We sub-selected the firing rate data from $t$=-250 ms to 250 ms and subtracted the baseline firing, estimated by averaging the firing rates from $t$=-250 ms to -150 ms for each trial. By estimating the firing rate for every time bin (=1 ms), our firing rate data became a ($N_{\text{trial}}$, $T$)-dimensional matrix for each neuron where $N_{\text{trial}}$ is a number of trials and $T$=501, the length of each trial. As for the kinematic parameters, peak velocity (PV) and duration were computed from an eye movement velocity profile for each trial. However, the distribution of saccade duration was skewed and can lead to inaccuracy in regression. In estimation, therefore, we used average velocity (AV), defined by 15°/(saccade duration), as a regression variable, instead of duration since the AV distribution was significantly more symmetric. Finally, we performed the multivariate linear regression of the firing rate data for the kinematic parameters (**Supplementary fig. 3a**) for each unit to find the model components for all unit data (**Supplementary fig. 3b,c**).

We checked the explanation power of the model fitted to each unit data, especially $\partial_z \mathbf{R}$, by computing the Akaike information criterion (AIC). We found that AIC decreased significantly ($P$<0.01, Student t-test) in all units when the PV-dependence is added ($\partial_{\mathrm{PV}} \mathbf{R}$) (MF: $\Delta$AIC=-65.33 $\pm$ 5.39, PC: -23.76 $\pm$ 1.65). Also, AIC significantly decreased ($P$<0.01, Student t-test) in a majority of the units (MF: $n$=99, PC: 83) when duration-dependence is augmented (MF: $\Delta$AIC=-23.60 $\pm$ 3.03, PC: -3.41 $\pm$ 0.48). Therefore, we confirmed that the model captured the true kinematic parameter-dependent trial-to-trial firing rate variability, not the data noise.

We also tested whether the models can predict the average firing rate profile. To do so, we split each data set into two, the training and test data. Then, we first constructed the rate models based on the training data sets. All the trials in the test data are split based on PV bins whose centers were 500°/s, 520°/s, ..., 660°/s and widths were 50°/s. We computed the PV-dependent average firing rate time series based on the estimated firing rates from the spike times of all the trials belonging to the PV bins. For each trial, we also computed the rate prediction from the training data and computed the prediction of the average firing rate in the same way as the test data (**Supplementary fig. 3d,f**). To test their agreement, we evaluated the peak firing rate and burst offset time as test measures for the average firing rates from the test data and model prediction (**Supplementary fig. 3e,g** ). This procedure was carried out for two different types of models; first, those parametrized only by PV (i.e., $\mathbf{z}$=[PV]) and the others with PV and duration (i.e., $\mathbf{z}$=[PV, AV=15°/(duration)]). We observed only an insignificant increase in the model performance by including duration as a parameter: With the PV-only model, $R^2$ for the peak firing rate versus PV were 0.929 $\pm$ 0.005 and 0.892 $\pm$ 0.023 for MFs and PCs, respectively. For the burst offset versus PV, $R^2$=0.887 $\pm$ 0.026 for MFs and 0.619 $\pm$ 0.095. With the PV-and-duration model, $R^2$ for the peak firing rate versus PV were 0.929 $\pm$ 0.005 for MFs and 0.791 $\pm$ 0.017 for PCs. For the burst offset versus PV, $R^2$=0.892 $\pm$ 0.021 for MFs and 0.702 $\pm$ 0.05 for PCs.

### Dimensionality Reduction by Principal Component Analysis with Perturbation

We developed a simple variant of principal component analysis (PCA) to perform dimensionality reduction of the population firing rate model. We first assume that, with any specific kinematic parameters (in the range of experimental observations), we can find a good dimensionally reduced representation by performing PCA on the population firing. Then, we find an approximation of the population firing and its change with kinematic parameters by another dynamical process with lower dimensionality. Our goal is that if we perform PCA on this approximated population firing, the result will be sufficiently close to that of the original population firing for any kinematic parameters. In our experimental data, the trial-to-trial variabilities of kinematic parameters and firing rate are relatively small. In this case, we can use the matrix perturbation theory[2,3] to find such an approximation of the population firing.

Our result can be summarized by Equation 2 in Materials and Methods: Given the kinematic parameter $\mathbf{z}$, if the time-dependent population firing $\mathbf{R}(t, \mathbf{z})$ of $N$ neurons is described by a linear model in Eq. S1, it can be approximated by the $K$-dimensional vectors $\mathbf{P}_K$ and $\partial_z \mathbf{P}_K$ ($K<N$) as

$$\begin{aligned} \mathbf{R}(t, \mathbf{z}) &\approx \mathbf{W}\left(\mathbf{P}_K + \sum_z \delta z\, \partial_z \mathbf{P}_K\right) + \mathbf{R}_\perp, \\ \partial_z \mathbf{P}_K &= \mathbf{W}^\dagger(\partial_z \mathbf{R}) + (\partial_z \mathbf{W})^\dagger(\mathbf{R}_0 - \mathbf{W}\mathbf{P}_K). \end{aligned} \tag{S2}$$

where $\mathbf{W}$ and $\partial_z \mathbf{W}$ are the weight matrices, as long as $\mathbf{R}_0$ admits $\mathbf{R}_0 \approx \mathbf{W}\mathbf{P}_K$ (**Supplementary fig. 6a,b** and **e,f**) and $\partial_z \mathbf{R}$ is sufficiently small. Furthermore, when we perform PCA on $\mathbf{R}$, the result is dominated by the first term since $\mathbf{R}_\perp$ makes a negligible contribution (see below). Therefore, we performed most of our manifold analysis in terms of $\mathbf{P}_K + \sum_z \delta z\, \partial_z \mathbf{P}_K$ (**Supplementary fig. 6c,d** and **g,h**), without considering $\mathbf{R}_\perp$. An exception is the dimensionally reduced MF firings given to the linear feed-forward network as inputs (Fig. 7). Here we use the full Eq. S2 for the approximate firing rates of individual neurons.

### Estimation of the model components

In the first step, we performed PCA on the kinematic-independent component, $\mathbf{R}_0$ (**Supplementary fig. 6a,b** and **e,f**). To do so, we computed the covariance matrix, $\mathbf{C} = \mathrm{Cov}[\mathbf{R}_0(t)]_t$ and its eigenvalues $\{\lambda_n\}$ ($\lambda_i \geq \lambda_j$ for $i > j$) with the corresponding eigenvectors $\mathbf{E} = [\mathbf{E}_1, \mathbf{E}_2, \dots, \mathbf{E}_N]$. If the first $K<N$ eigenvalues are dominant (see below for the determination of $K$), a dimensionally reduced approximation of $\mathbf{R}_0$ can be obtained by the projection of the population activity to a $K$-dimensional subspace of $\mathbf{E}$ as

$$\mathbf{R}_0 \approx \mathbf{W}\mathbf{P}_K, \quad \mathbf{P}_K = \mathbf{W}^\dagger \mathbf{R}_0, \quad \mathbf{W} = [\mathbf{E}_1, \dots, \mathbf{E}_K].$$

Then, in the second step, we approximated the full covariance matrix of $\mathbf{R}(t, \mathbf{z})$ as

$$\hat{\mathbf{C}} = \mathrm{Cov}[\mathbf{R}(t, \mathbf{z})]_t \approx \mathbf{C} + \sum_z \delta z\, \partial_z \mathbf{C}, \quad \partial_z \mathbf{C} = \mathrm{Cov}[\mathbf{R}_0, \partial_z \mathbf{R}]_t + \mathrm{Cov}[\partial_z \mathbf{R}, \mathbf{R}_0]_t,$$

assuming that the kinematics-dependent part is sufficiently small. When the second $\delta z$-dependent terms are small, they can be considered as small perturbations. In that case, the Rayleigh–Schrödinger perturbation theory tells that eigenvalues $\{\hat{\lambda}_n\}$ and eigenvectors $\hat{\mathbf{E}} = [\hat{\mathbf{E}}_1, \dots, \hat{\mathbf{E}}_N]$ of $\hat{\mathbf{C}}$ are approximately[2,3],

$$\begin{aligned} \hat{\lambda}_n &\approx \lambda_n + \sum_z \delta z \, \partial_z \lambda_n, \quad \partial_z \lambda_n = \mathbf{E}_n^\dagger (\partial_z \mathbf{C}) \mathbf{E}_n, \\ \hat{\mathbf{E}}_n &\approx \mathbf{E}_n + \sum_z \delta z \, \partial_z \mathbf{E}_n, \quad \partial_z \mathbf{E}_n = \sum_{k \neq n} \frac{\partial_z \lambda_n}{\lambda_n - \lambda_k} \mathbf{E}_k. \end{aligned} \tag{S3}$$

**Supplementary fig. 6i** shows an example of the PC data. The eigenvalues of the covariance matrix change is well approximated by Eq. S3. **Supplementary fig. 6j** shows how much contribution each $\partial_z \mathbf{E}_i$ gets from $\mathbf{E}_j$. We can see that $\partial_z \mathbf{E}_3$ and $\partial_z \mathbf{E}_4$ especially get significant contributions not only from the first four $\mathbf{E}_j$'s but also from the higher ($K$>4) dimensional components.

Finally, by using the eigenvector perturbation in Eq. S3, we determined the rest of the components in Eq. S2,

$$\begin{aligned} \partial_z \mathbf{W} &= [\partial_z \mathbf{E}_1, \partial_z \mathbf{E}_2, \dots, \partial_z \mathbf{E}_K], \\ \mathbf{R}_\perp &= -\sum_z \delta z \, \mathbf{E}_\perp (\partial_z \mathbf{E}_\perp)^\dagger \mathbf{W} \mathbf{P}_K, \end{aligned} \tag{S4}$$

where $\mathbf{E}_\perp = [\mathbf{E}_{K+1}, \dots, \mathbf{E}_N]$, and $\partial_z \mathbf{E}_\perp = [\partial_z \mathbf{E}_{K+1}, \partial_z \mathbf{E}_{K+2}, \dots, \partial_z \mathbf{E}_N]$. We give the detailed derivation in the next section. Therefore, $\partial_z \mathbf{W}$ represents a contribution from the eigenvector perturbation, and we will call it an *indirect projection*, while we will call $\mathbf{W}^\dagger (\partial_z \mathbf{R})$ a *direct projection* term. On the other hand, $\mathbf{R}_\perp$ represents the $K$-dimensional approximation of $\mathbf{R}_0$ rotating out of the $K$-subspace by perturbation and makes a negligible contribution when we perform PCA. **Supplementary fig. 6k,l** show that we can well predict the PCA results of the population firing given the changes in a kinematic parameter, PV, without $\mathbf{R}_\perp$, while the indirect projection part can make a substantial contribution.

### Determination of $K$

As a final note, we explain how we determined $K$ in the first step: we first found $K$ components that explained more than 85% of the total variance in $\mathbf{R}_0$. This criterion gave us $K = 2$ and 4 for MFs and PCs, respectively. We also computed the participation ratio[4,5], $(\sum_n \lambda_n)^2 / \sum_n \lambda_n^2$, which estimated $K$=2 (MFs) and 3 (PCs), from $\mathbf{R}_0$. However, $K$=4 for PCs was more robust when we varied kinematic parameters or hyperparameters such as the smoothing time scale for rate estimation. Finally, the cross-validation analysis for PCA[6] of $\mathbf{R}_0$ also confirmed $K = 2$ and

4: We first randomly selected 70% of elements in $\mathbf{R}_0$ matrix ("test data") and replaced them by Gaussian random numbers, leaving the other 30% of the "training data" untouched. Using the data with random replacements, we repeatedly performed PCA until we got the stable prediction of the test data. Then, we computed the cross-validation error by the squared sum of the differences between the predicted and real test data. This procedure was repeated 200 times for each $K$ from 2 to 20. We found that the cross-validation error, averaged over the repetitions, was minimal at $K$=2 and 4 for the MF and PC data, respectively.

### Derivation of Eq. S2 and S4

Given the perturbed eigenvectors in Eq. S3, the projection of the population activity to them, $\hat{\mathbf{P}}$, is

$$\hat{\mathbf{P}} = \hat{\mathbf{E}}^\dagger \mathbf{R} \approx \mathbf{P} + \sum_z \delta z \left\{ \mathbf{E}^\dagger (\partial_z \mathbf{R}) + (\partial_z \mathbf{E})^\dagger \mathbf{R}_0 \right\}$$

where $\mathbf{E} = [\mathbf{E}_1, \ldots, \mathbf{E}_N]$ and $\partial_z \mathbf{E} = [\partial_z \mathbf{E}_1, \ldots, \partial_z \mathbf{E}_N]$. If the left inverse of $\hat{\mathbf{E}}^\dagger$ is $\hat{\mathbf{U}} = \mathbf{U} + \sum_z \delta z \partial_z \mathbf{U} + O(\delta z^2)$, we get $\mathbf{U} = \mathbf{E}$ and $\partial_z \mathbf{U} = -\mathbf{E}(\partial_z \mathbf{E})^\dagger \mathbf{E}$, from the condition $\hat{\mathbf{U}}\hat{\mathbf{E}}^\dagger = \mathbf{1}$.

Now we find the low dimensional representation $\hat{\mathbf{P}}_K$ by keeping only the first $K$ components of $\hat{\mathbf{P}}$, i.e. $\hat{\mathbf{P}}_K = (\hat{\mathbf{P}})_K$ where $(\cdot)_K$ denotes selecting only the first $K$ rows in a matrix. With $\hat{\mathbf{W}} = [\hat{\mathbf{U}}_1, \hat{\mathbf{U}}_2, \ldots, \hat{\mathbf{U}}_K]$ and again $\mathbf{W} = [\mathbf{E}_1, \mathbf{E}_2, \ldots, \mathbf{E}_K]$,

$$\mathbf{R} \approx \hat{\mathbf{W}}\hat{\mathbf{P}}_K = \mathbf{W}\mathbf{P}_K + \sum_z \delta z \left\{ \mathbf{W}(\partial_z \mathbf{P})_K + (\partial_z \mathbf{U}_\parallel)\mathbf{P}_K \right\} + O(\delta z^2)$$

where $\partial_z \mathbf{U}_\parallel = [\partial_z \mathbf{U}_1, \partial_z \mathbf{U}_1, \ldots, \partial_z \mathbf{U}_K]$.

Through a little algebra, this equation can be rewritten as

$$\begin{aligned} \mathbf{R} \approx \mathbf{W}\mathbf{P}_K &+ \sum_z \delta z \mathbf{W}\mathbf{W}^\dagger (\partial_z \mathbf{R}) + \sum_z \delta z \mathbf{W}(\partial_z \mathbf{W})^\dagger (\mathbf{R}_0 - \mathbf{W}\mathbf{P}_K) \\ &- \sum_z \delta z \mathbf{E}_\perp (\partial_z \mathbf{E}_\perp)^\dagger \mathbf{W}\mathbf{P}_K \end{aligned}$$

where $\partial_z \mathbf{W} = [\partial_z \mathbf{E}_1, \partial_z \mathbf{E}_2, \ldots, \partial_z \mathbf{E}_K]$. $\mathbf{E}_\perp$ and $\partial_z \mathbf{E}_\perp$ are both orthogonal to the $K$-subspace as $\mathbf{E}_\perp = [\mathbf{E}_{K+1}, \ldots, \mathbf{E}_N]$, and $\partial_z \mathbf{E}_\perp = [\partial_z \mathbf{E}_{K+1}, \partial_z \mathbf{E}_{K+2}, \ldots, \partial_z \mathbf{E}_N]$. Defining $\mathbf{R}_\perp = -\sum_z \delta z \mathbf{E}_\perp (\partial_z \mathbf{E}_\perp)^\dagger \mathbf{W}\mathbf{P}_K$, we reach Eq. S2.

### Alignment of Two Manifolds by Canonical Correlation Analysis

In the main text, we compared manifolds obtained from two or more different data sets, such as firings recorded with the left-directed saccades and those with the right-directed ones. We used canonical correlation analysis (CCA) to verify if a pair of manifolds can be related by a linear transformation[7–9]. We first used the CCA for the kinematics-independent components of two data sets to find the best linear alignment transformation between them. Then, we transformed the kinematics-dependent parts by the alignment transform and checked how well they match with each other.

If there are two data called $A$ and $B$, we denote the kinematics-independent components ($\mathbf{P}_K$ in Eq. S2) of their manifolds, $\mathbf{P}_A$ and $\mathbf{P}_B$, respectively. Also, we discretize time and consider them as $N \times T$ matrices instead of time-dependent vectors, where $T$ is the length of trials.

First, we perform the QR decomposition,

$$\mathbf{P}_X^\dagger = \mathbf{Q}_X \mathbf{V}_X, \quad X = A, B.$$

From the singular value decomposition of the comparison matrix $\mathbf{Q}_A^\dagger \mathbf{Q}_B = \mathbf{U}_A \mathbf{S} \mathbf{U}_B^\dagger$, we obtain the transformation matrices to the best aligned manifolds, $\tilde{\mathbf{P}}_A$ and $\tilde{\mathbf{P}}_B$, as

$$\tilde{\mathbf{P}}_X^\dagger = \mathbf{P}_X^\dagger \mathbf{M}_X, \quad \mathbf{M}_A = \mathbf{V}_X^{-1} \mathbf{U}_X, \quad X = A, B.$$

The diagonal elements of $\mathbf{S}$ are correlations between $\tilde{\mathbf{P}}_A$ and $\tilde{\mathbf{P}}_B$. We obtain a representation of $\mathbf{P}_B$ aligned to $A$

$$\mathbf{P}_{B\to A}^\dagger = \mathbf{P}_B^\dagger \mathbf{T}_{B\to A}, \quad \mathbf{T}_{B\to A} = \mathbf{M}_B \mathbf{M}_A^{-1}/N,$$

where $N$ is a norm of $\mathbf{M}_B \mathbf{M}_A^{-1}$ to maintain the size difference between two manifolds. Then, Eq. S2 becomes

$$\begin{aligned} \hat{\mathbf{R}}_B &\approx \mathbf{W}_B \left( \mathbf{P}_B + \sum_z \delta z \, \partial_z \mathbf{P}_B \right) + \mathbf{R}_\perp \\ &= \mathbf{W}_B \mathbf{T}_{A\to B}^\dagger \mathbf{P}_{B\to A} + \sum_z \delta z \, \mathbf{W}_B \mathbf{T}_{A\to B}^\dagger \mathbf{T}_{B\to A}^\dagger \partial_z \mathbf{P}_B + \mathbf{R}_\perp \\ &= \mathbf{W}_{B\to A} \left\{ \mathbf{P}_{B\to A} + \sum_z \delta z \, \partial_z \mathbf{P}_{B\to A} \right\} + \mathbf{R}_\perp, \end{aligned} \tag{S5}$$

where $\mathbf{W}_{B\to A} = \mathbf{W}_B \mathbf{T}_{A\to B}^\dagger$ is a new weight matrix for $\mathbf{P}_{B\to A}$ and $\partial_z \mathbf{P}_{B\to A} = \mathbf{T}_{B\to A}^\dagger (\partial_z \mathbf{P}_B)$ is the aligned manifold perturbation.

#### Linear Feed-forward Network (LFFN) Models

In the main text, we considered three different types of LFFN. They commonly had the movement kinematics-independent and dependent components for output variables ($\mathbf{Y}$, $\partial_{\mathbf{z}}\mathbf{Y}$) and input ($\mathbf{X}$, $\partial_{\mathbf{z}}\mathbf{X}$). Then, we compute the expectation value of the least-square error given the distribution of $\mathbf{z}$, $p(\mathbf{z})$,

$$
\begin{aligned}
E(\mathbf{T}) &= \int d\mathbf{z}\, p(\mathbf{z}) \left\| \mathbf{Y} + \sum_{z} \delta z\, \partial_z \mathbf{Y} - \mathbf{T}\left( \mathbf{X} + \sum_{z} \delta z\, \partial_z \mathbf{X} \right) \right\|^2 \\
&= \int d\mathbf{z}\, p(\mathbf{z}) \left\| \mathbf{Y} - \mathbf{TX} \right\|^2 + \int d\mathbf{z}\, p(\mathbf{z}) \sum_{z} \delta z \left( \partial_z \mathbf{Y} \right) \cdot \mathbf{TX} + (\mathbf{X} \leftrightarrow \mathbf{Y}) \\
&\quad + \int d\mathbf{z}\, p(\mathbf{z}) \sum_{z} \sum_{z'} \delta z \delta z' \left( \partial_z \mathbf{Y} - \mathbf{T}\partial_z \mathbf{X} \right) \cdot \left( \partial_{z'} \mathbf{Y} - \mathbf{T}\partial_{z'} \mathbf{X} \right).
\end{aligned}
$$

We assume that $p(\mathbf{z})$ is approximately the Gaussian distribution with zero mean. Then, we get the error function in Equation 3 in the main text,

$$
E(\mathbf{T}) = \left\| \mathbf{Y} - \mathbf{TX} \right\|^2 + \sum_{z} \sum_{z'} \mathrm{Cov}[z, z'] \left( \partial_z \mathbf{Y} - \mathbf{T}\partial_z \mathbf{X} \right) \cdot \left( \partial_{z'} \mathbf{Y} - \mathbf{T}\partial_{z'} \mathbf{X} \right). \tag{S6}
$$

In evaluating the performance, we measured the total variability by replacing the prediction ($\mathbf{TX}$, $\mathbf{T}\partial_z\mathbf{X}$) in Eq. S6 by the mean $\langle \mathbf{Y} \rangle$ and $\langle \partial_z \mathbf{Y} \rangle$ and compared it to $E(\mathbf{T})$ to evaluate the goodness of fit, $R^2$. To prevent overfitting, we used the LASSO scheme[10] to minimize $E(\mathbf{T}) + \sum_{i,j} \lambda_i \left| T_{ij} \right|$. We used the MATLAB package *glmnet*[11] and chose optimal $\lambda_i$ for each $Y_i$ by finding where AIC minimizes.

The communication subspace model (**Supplementary fig. 8d-e**) was obtained by the rank-reduced regression[12] with the error function in Eq. S6. We first performed the unconstrained optimization of $E(\mathbf{T})$ to find the optimal least-square solution $\mathbf{T}_{\mathrm{OLS}}$. Then, we computed the $d$ principal components of the predictor, $\mathbf{V}$, and obtained the reduced-rank solution $\mathbf{T}_{\mathrm{RRR}} = \mathbf{V}\mathbf{V}^{\dagger}\mathbf{T}_{\mathrm{OLS}}$.
